# STopover captures spatial colocalization and interaction in the tumor microenvironment using topological analysis in spatial transcriptomics data

**DOI:** 10.1101/2022.11.16.516708

**Authors:** Sungwoo Bae, Hyekyoung Lee, Kwon Joong Na, Dong Soo Lee, Hongyoon Choi, Young Tae Kim

## Abstract

Unraveling the spatial configuration of the tumor microenvironment (TME) is key to understanding tumor-immune interactions to translate them into immuno-oncology. With the advent of spatially resolved transcriptomics (SRT), the TME could be dissected for whole cell types across numerous RNAs. We suggest a novel approach, STopover, which performs topological analysis to compute the colocalization patterns between cell types and map the location of cell□cell interactions. While gradually lowering the threshold for the feature, the connected components (CCs) were extracted based on the spatial distance between the unit tissue region and the persistence of the CCs. Local and global Jaccard indices were calculated between the CCs of a feature pair to measure the extent of spatial overlap. The STopover was applied to various lung cancer data obtained from SRT platforms, both barcode and image-based SRT, and could explain the infiltration patterns of immune and stromal cells in the TME. Moreover, the method predicted the top cell□cell communication based on the ligand□receptor database and highlighted the main region of the interaction. STopover is a tool to decipher spatial interaction in the tissue and shed light on the pathophysiology underlying the microenvironment.

## Main

One of the essential characteristics of cancer progression is acquiring an immune escape mechanism in the tumor microenvironment (TME)^1^. The heterogeneous cells in the TME constitute the landscape to evade the immune system, which has been regarded as a target for immunotherapy^2^. The most widely investigated mechanism is augmenting the immune checkpoint, which was designed to protect normal cells from cytotoxic immune cells. Enhancement of either axis of checkpoints eventually leads to impaired function or apoptosis of cytotoxic immune cells such as CD8+ T cells^3, 4^. Alternatively, MHC class I expression, which is responsible for the antigen presentation of cancer cells, is reduced^5^, or cancer interacts with cancer-associated fibroblasts (CAFs) and alters the microenvironment such that immune cells have more difficulty suppressing tumor growth^6^. Recently, interest has been growing in the field of cancer immunotherapy, particularly targeting PD-1, PD-L1, and CTLA-4, which have been widely investigated in clinical research. Multiple preclinical studies and clinical trials have demonstrated that immunotherapy targeting various mechanisms in the TME combined with conventional chemotherapy results in effective tumor suppression.^7^.

However, not all patients respond well to immunotherapy, even though predictive biomarkers for immune checkpoint inhibitors (ICIs), such as PD-L1 expression, microsatellite instability, and tumor mutational burden, have been used in the clinic^4^. As a spatial feature of the TME related to the immunotherapy response, the infiltration pattern of immune cells in the tumor is regarded as a key to characterizing immune status^8^. In the case of an immune-excluded or immune-desert tumor where immune cells do not penetrate the tumor, the response to immunotherapy is bound to be poor. Moreover, not only tumor and immune cells but also stromal components in the tumor microenvironment (TME) influence immune cell action on cancer cells^9^. Given that the therapeutic response of ICI is determined by complex intercellular interactions, including lymphocytes, myeloid cells, and fibroblasts in the TME, a platform that simultaneously analyzes the spatial configuration of multiple cell types is needed.

With the introduction of spatially resolved transcriptomics (SRT), which acquires the location of RNAs and their quantitative expression in the tissue, the spatial characteristics of cells can be comprehensively analyzed^10^. Because this technique provides whole gene expression data with spatial information, it has the potential to comprehensively analyze the spatial configuration and complex interactions of various cells in the TME. Here, we propose a method, STopover, which applies topological analysis to SRT to extract colocalization patterns between cell types in cancer tissue and to predict the degree of intercellular interaction. Our suggested new approach provides the extent of topological overlap between various cells, such as tumor and immune cells, and presents an estimate of the spatial interaction between cells. STopover can provide quantitative information on spatial interactions in the TME beyond spatial gene expression or cell enrichment.

## Results

### STopover captures colocalized regions of a feature pair in a simulation dataset

The STopover was designed to quantify the topological colocalization of two given features using Morse filtration (**Fig. 1 and Supplementary Fig. 1**). The threshold of cell abundance or gene expression (features) observed at each tissue region was gradually lowered from the highest value to the lowest value to find connected components (CCs) based on spatial proximity. Next, it captures the locally activated region of features by reconfiguring the CCs based on the minimum size of the CC and the threshold range that a certain CC is continuously observed^11^. After removing CCs with low average feature values, the degree of local and global spatial overlap between the two features was computed by the Jaccard index. Finally, by applying STopover to the barcode- and image-based SRT, it was possible to visualize the extent of overlap between tumor cells, immune cells, and fibroblasts and to present an estimate of the spatial interaction between cells. First, the utility of STopover, which extracts topographically overlapping tissue domains between the cell type pairs, was examined in a simulation dataset and compared with the conventional threshold-based approach. As an example of the threshold-based approach to delineating the overlapping tissue regions of the two cell types, a threshold filter that removes the value below 20% of the maximum was applied to segment the main tissue domain where the two cell types are colocalized. The threshold-based method could capture cancer and immune cell overlap in the region where both cell types showed high abundance (Region A in **Fig. 2a, b**). However, that method could not accurately delineate the key location where cancer cells are scarce while immune cells are highly abundant (Region B in **Fig. 2a, b**). Thus, the conventional method can easily miss the locally active regions of cancer and immune interactions. In contrast, STopover could precisely capture the colocalized tissue domains between cancer and immune cells (Region B **in Fig. 2a, c**). Moreover, as quantitative information, STopover could rank the degree of overlap between the CCs of cancer and immune cells with the Jaccard index and indicate subregions where high cancer-immune interactions are expected (**Fig. 2d**).

**Fig. 1:**
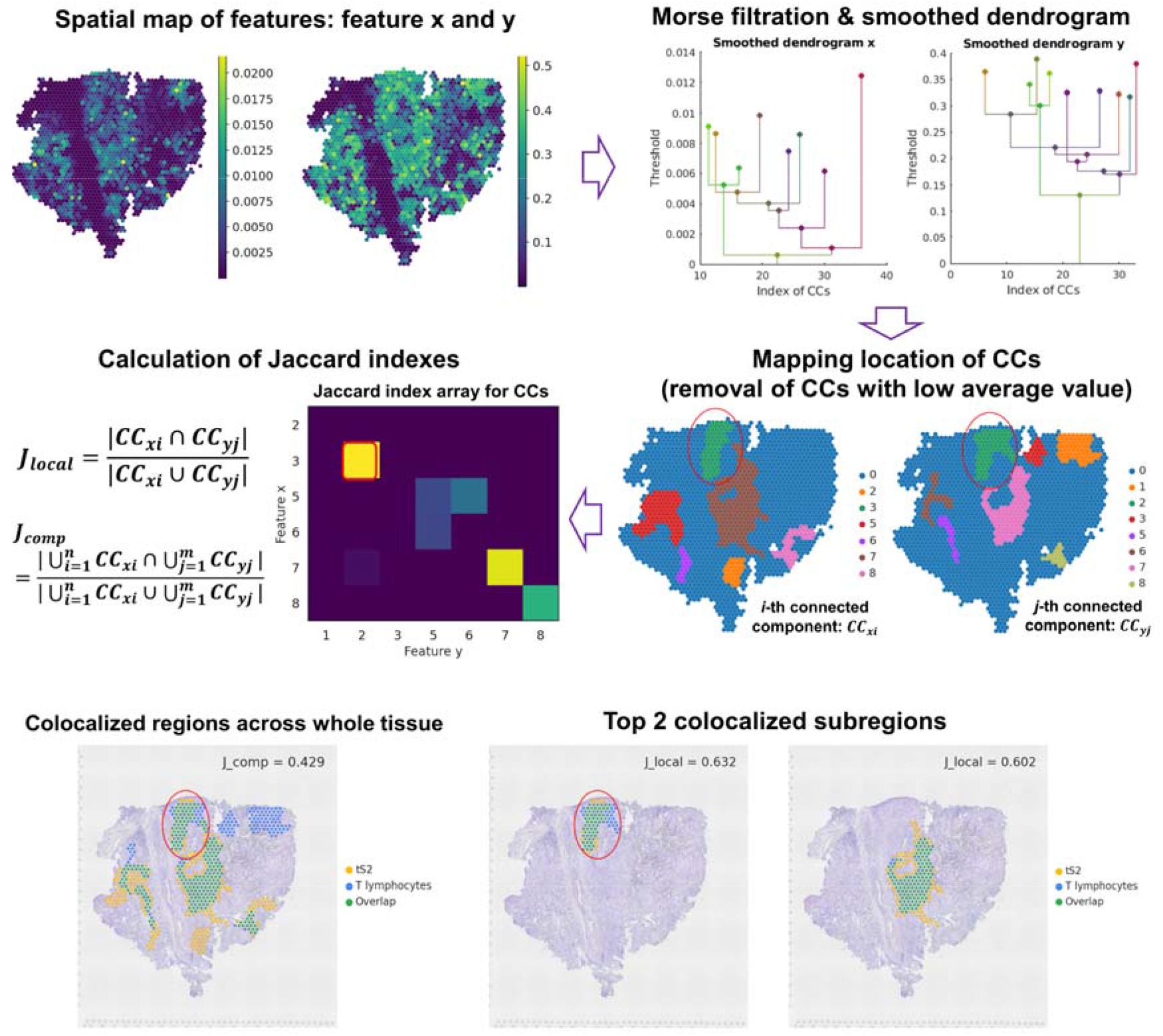
Schematic image of STopover. A STopover is a tool that utilizes spatially resolved transcriptomics (SRT) and applies topological analysis to extract colocalization patterns between the cell types and estimate the spatial cell□ cell interaction in the tumor microenvironment. The spatial distribution of cell types is given as inputs to the STopover model when analyzing cell□cell colocalization. The LR pairs from the CellTalkDB database^15^ are provided as inputs when estimating spatial cello cell communication mediated by ligand receptor (LR) interactions. By utilizing Morse filtration and dendrogram smoothing processes, the key locations of the overlapping spatial domain were extracted as connected components (CCs). After removing CCs with low average feature values, the Jaccard indices were calculated for every CC pair between the two features and named *J_local_*. The CC pairs with a large Jaccard index indicate important tissue subregions where the two features are highly colocalized. Additionally, all CCs from each feature were aggregated, and the Jaccard index between the two aggregated CCs was calculated and named *J_comp_*. *J_comp_* measures the extent of spatial overlap of the two features on the global scale.

**Fig. 2:**
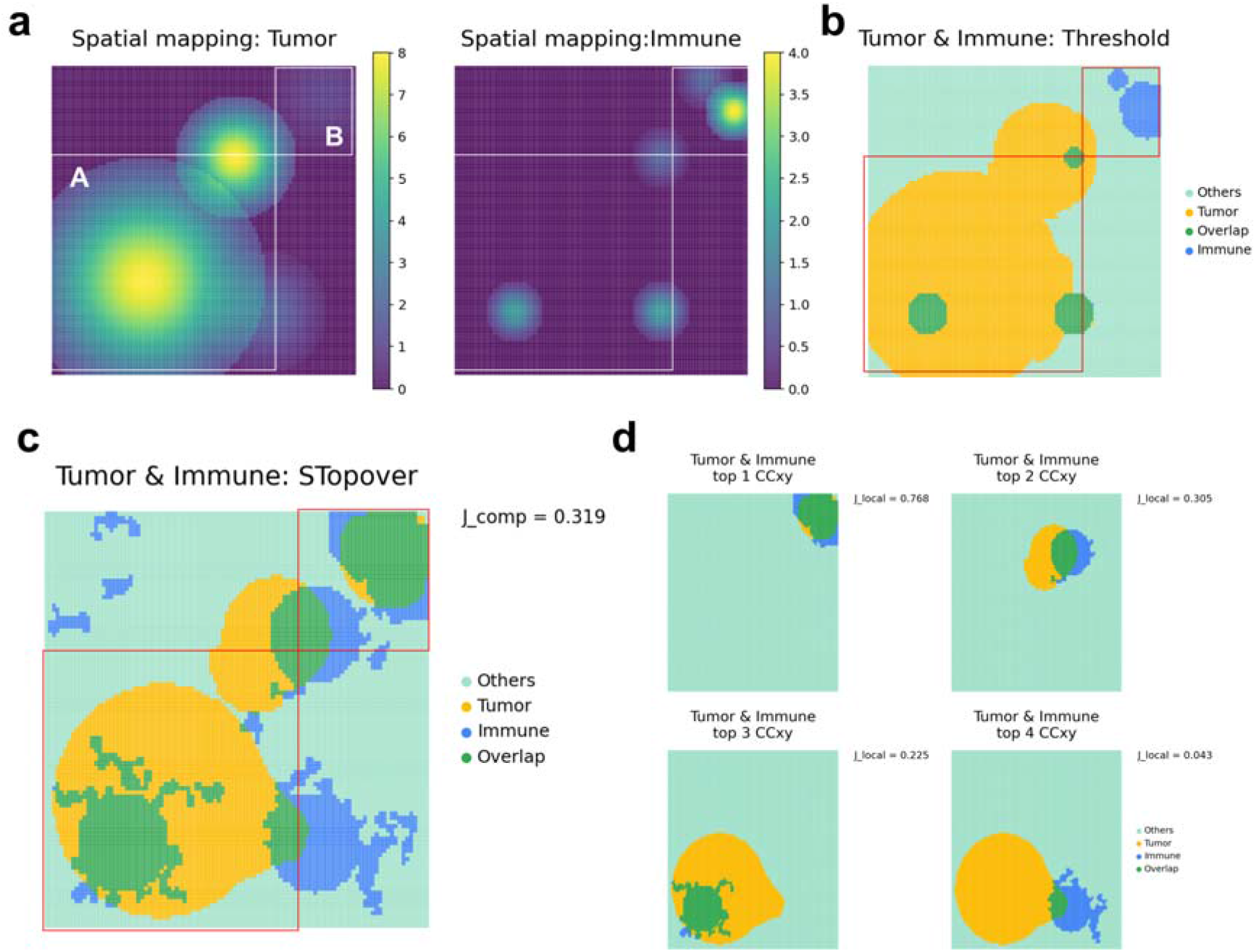
STopover reveals a subregional colocalization pattern in the simulation dataset. A simulation dataset was created to examine the ability of STopover to capture small but highly colocalized subregions. Trimmed 2D Gaussian functions were applied multiple times to a 100 by 100 grid, and the activity of tumor and immune cells was simulated. **a**, The spatial maps of tumor and immune cell activity were visualized with a colormap. **b**, A threshold-based approach was applied, and the regions where the activity was above 20% of the maximum were filtered to segment the key regions of tumor (yellow) and immune cells (blue). Then, spatially overlapping domains between the two cell types are highlighted in green. **c**, STopover was applied to segment the main patterns of tumor (yellow) and immune cell (blue) activity as CCs. The CCs for the tumor (yellow) and immune cells (blue) and the intersecting subregions (green) between the two aggregated CCs were visualized on the grid. **d**, The top 4 CC pairs between tumor and immune cells showing the highest spatial overlap represented by the *J_local_* score were visualized. The CC from the tumor (yellow), immune cells (blue), and intersecting subregions (green) was visualized on the grid. Overall, STopover was superior to the conventional threshold-based method in capturing locally active subregions where tumor cell activity is low but immune cell activity is high (Region B).

### STopover reveals spatial overlap patterns between lung cancer cell types in barcodebased SRT

To assess the usefulness of the method in cancer tissues, STopover was applied to the human lung adenocarcinoma (ADC) Visium datasets. Lung cancer is one of the cancer types in which immunotherapy plays a crucial role in patient treatment^12^. First, reference single-cell data obtained from human lung cancer tissue^13^ were integrated with spatial data using CellDART^14^ to estimate the spatial distribution of the major cell types. Then, the localization patterns of the cancer cell type (‘tS2’), which is related to lung ADC progression^13^, and other main cell types (‘Fibroblasts’, ‘Endothelial cells’, ‘Myeloid cells’, ‘MAST cells’, ‘NK cells’, ‘B lymphocytes’, and ‘T lymphocytes’) were extracted by STopover. The composite overlap score was calculated by measuring the Jaccard index between the combined CC aggregates between the two cell types (*J_comp_*) (**Fig. 1**). In addition, the local overlap score (*J_local_*) was computed by measuring the Jaccard index between every CC pair of the two cell types. *J_comp_* represents the degree of global overlap between the two cell types, while *J_local_* shows tissue subregions where the cell distribution is highly colocalized.

Two representative slides were selected among 11 tissue sections to evaluate whether STopover accurately extracts the spatial overlap patterns of cells in the TME. One of the tissues (‘*spa06ca01*’) had low PD-L1 expression (0%) and showed immune-excluded patterns by visual inspection (**Supplementary Fig. 2a**). The other tissue (‘*spa18ca02*’) had high PD-L1 expression (100%) and presented immune-inflamed patterns (**Supplementary Fig. 2b**). When the spatial distribution of cell types was extracted as CCs and mapped to the tissues, the CC distribution was concordant with spatial cell type patterns (**Supplementary Fig. 2 and Fig. 3**). The spatial overlap pattern between cell types in two lung ADC samples exhibited disparate patterns of tumor, immune, and stromal distribution (**Fig. 3**). In the PD-L1 low tissue, the spatial patterns of tS2 and immune cells, including MAST, NK, and T cells, were highly exclusive (**Fig. 3a**). Accordingly, the immune cell subtypes were among the top 3 cell types with the lowest *J_comp_* values. In contrast, in the PD-L1 high tissue, the patterns of tS2 and MAST, NK, or T cells overlapped in several tissue subregions (**Fig. 3b**), with *J_local_* ranging from 0.108 to 0.848 in MAST cells, 0.025 to 0.515 in NK cells, and 0.024 to 0.632 in T cells (**Supplementary Fig. 3**). The tissue subregions with the top *J_local_* scores (**Supplementary Fig. 3**) were shared between the three immune cell types. Additionally, MAST, NK, and T cells belonged to the top 3 cell types with the highest *J_comp_* (**Fig. 3b**). Notably, in the PD-L1 low tissue, fibroblasts colocalized with tS2 at the border of the tumor defined by CCs of tS2; however, in PD-L1 high tissue, the two cell types showed little overlap.

**Fig. 3.**
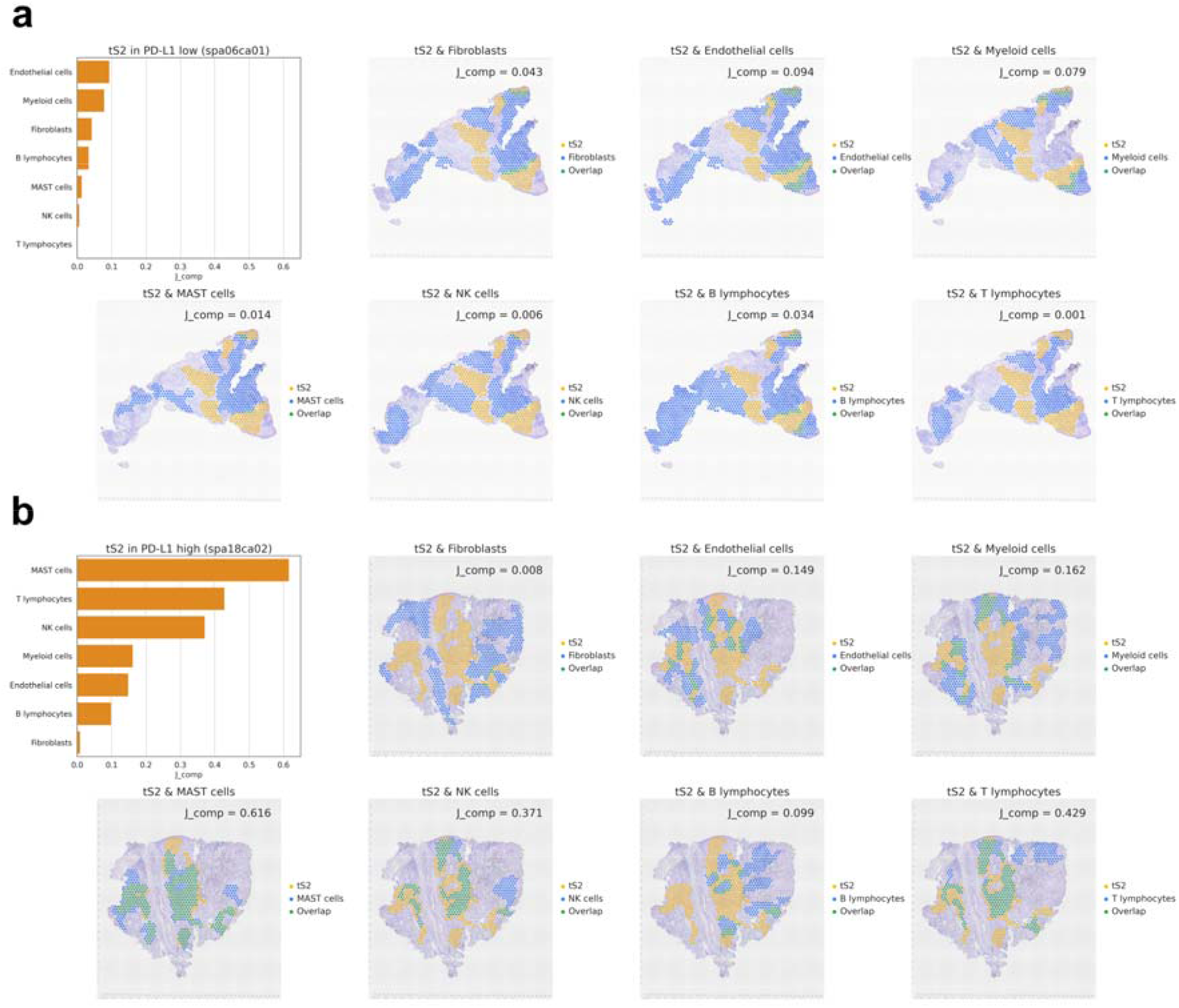
STopover explains the spatial configuration of the TME in lung cancer tissues using barcode-based SRT. STopover was applied to barcode-based SRT of lung cancer tissues with high PD-L1 expression (*spa06ca01*, 0%) and low PD-L1 expression (*spa18ca02*, 100%). The spatial colocalization patterns between tS2, one of the cancer epithelial subtypes associated with the progression of cancer, and other main cell types (fibroblasts, endothelial cells, myeloid cells, MAST cells, NK cells, B lymphocytes, and T lymphocytes) were investigated in both tissues. **a,** The *J_comp_* values between tS2 and the other cell types in spa06ca01 tissue were visualized as a bar plot in the top left corner of the plot. Additionally, the aggregated CCs for tS2 (yellow) and other main cell types (blue) were mapped to the tissue, and the intersecting tissue domain was highlighted in green. **b,** *J_comp_* values for tS2 and other cell types in spa18ca02 tissue were also visualized with a barplot, and the CC locations were mapped to the tissue. The two selected PD-L1 high and low tissues showed converse patterns of cell infiltration in the tumor, and the extent of infiltration could be measured as *J_comp_*.

To further dissect the tumor and T cell interactions in PD-L1 high and low tumors, the spatial relationship between cancer cells and T cell subtypes was investigated. Spatial overlap patterns of tS2 with T cell subtypes, including naïve CD4+ T cells, CD4+ helper T cells (Th), CD8+/CD4+ mixed Th cells, exhausted follicular Th (Tfh) cells, regulatory T cells (Treg), naïve CD8+ T cells, cytotoxic CD8+ T cells, exhausted CD8+ T cells, CD8low T cells, and other nonspecified T cells (T lymphocytes_ns), were calculated. Overall, PD-L1 low tumors did not show significant spatial overlap between various T cell subtypes and tS2 (*J_comp_*: 0.000–0.142), except for exhausted T cell subtypes, which were colocalized at the border of tS2 (**Supplementary Fig. 4a**). Conversely, PD-L1 high tumor, naïve, effector, and exhausted T cell subtypes were highly colocalized with tS2 (*J_comp_*: 0.318–0.530) (**Supplementary Fig. 4b**). Interestingly, Tregs showed the lowest overlap with tS2 (*J_comp_*: 0.101), while cytotoxic CD8+ T cells presented the highest overlap with tS2 (*J_comp_*: 0.530). This finding implies that the TME is less suppressive to T lymphocytes in the selected PD-L1 high tissue (*spa18ca02*)compared to the PD-L1 low tissue (*spa06ca02*).

Then, STopover analysis was extensively performed across 11 lung cancer Visium datasets, and the spatial configuration of the TME in various lung cancer tissues was explored. A total of 6 tissues (*spa01ca01, spa01ca02, spa02ca01, pa02ca02, spa06ca01, spa10ca01*) had low PD-L1 expression (0% in all samples), and the other 6 tissues (*spa16ca01, spa17ca01, spa17ca02, spa18ca01, spa18ca02*) had high PD-L1 expression (range: 80-100) (**Supplementary Table 1**). First, the correlation between pseudobulk expression of ICI response biomarkers (PD-L1 and MHC class I) and spatial overlap of tS2 and T cells represented by *J_comp_* as a quantitative value for representing topological T cell infiltration was examined. The expression of *HLA-B* and *HLA-C*, which encode MHC class I, showed a significant positive correlation with *J_comp_*; however, *CD274*, which encodes PD-L1, and *PDCD1*, which encodes PD-1, did not show a significant correlation (**Supplementary Fig. 5**). Second, *J_comp_* was calculated between tS2 and other main cell types and was visualized with a heatmap (**Fig. 4a**). Among the 7-cell types, MAST, NK, and T cells showed a similar trend of *J_comp_* values across the tissue samples and were clustered together. Meanwhile, the 11 tissue samples were clustered into two groups (‘Cluster 1’ and ‘Cluster 2’) according to the trend of spatial overlap between tS2 and other cell types. Overall, compared to Cluster 1, Cluster 2 showed higher median *J_comp_* scores in MAST, NK, B, and T cells and lower scores in fibroblasts, endothelial cells, and myeloid cells (**Fig. 4b**). Thus, Cluster 2 represents the TME with high infiltration of lymphocytes and low involvement of myeloid cells and fibroblasts. In summary, STopover can describe and quantify the spatial relationship between cancer-immune and cancer-stromal components in lung cancer, particularly in barcode-based SRT.

**Fig. 4.**
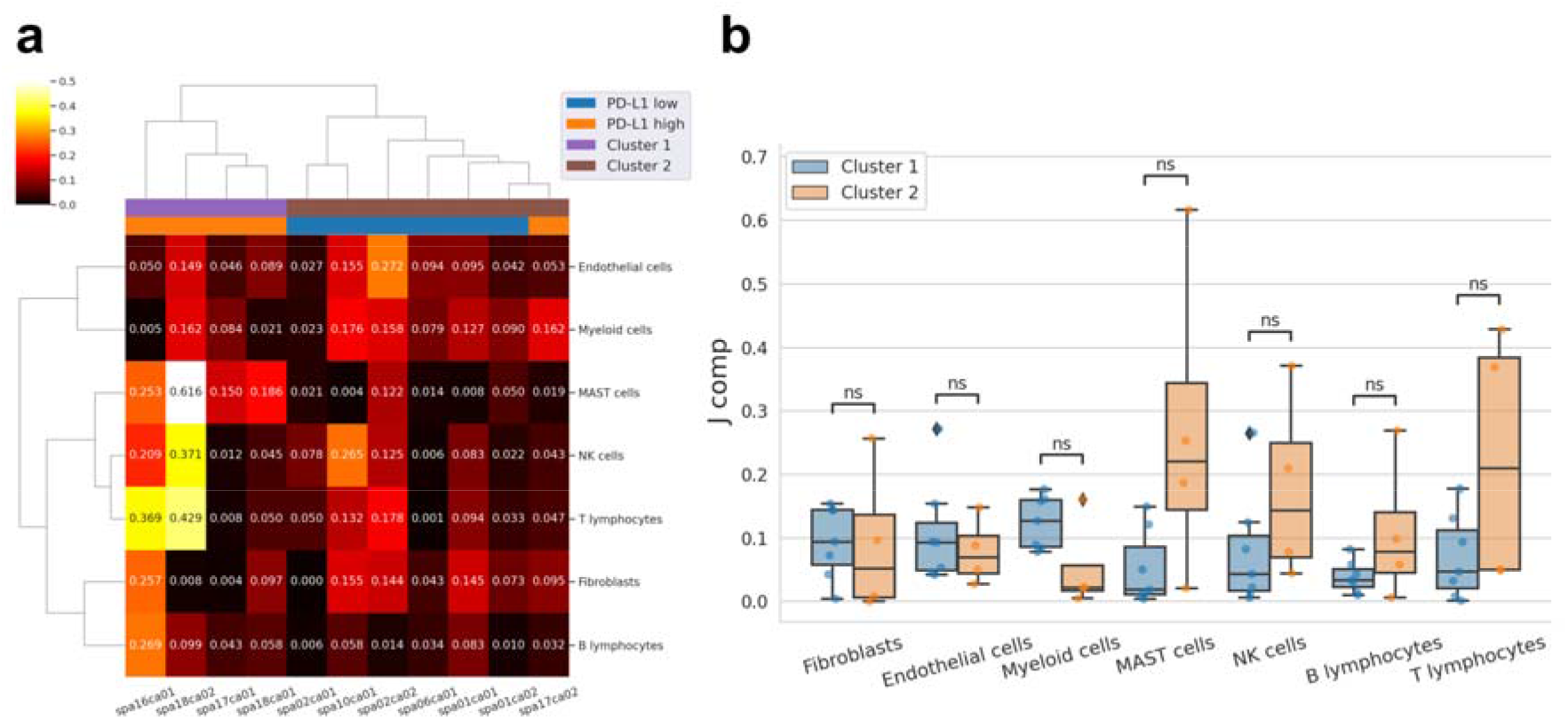
STopover clusters multiple lung cancer tissues based on the cell colocalization pattern in the TME using barcode-based SRT. The barcode-based SRTs of 11 lung cancer tissues were analyzed with STopover. The extent of spatial overlap between tS2 and other main cell types was represented by *J_comp_* scores. **a,** *J_comp_* values between tS2 and other main cell types in 11 lung cancer tissues were visualized with a heatmap. The Pearson correlation distances were computed across all cell type pairs and all tissue pairs, and hierarchical clustering was performed. The tissues were classified into two clusters: Clusters 1 and 2. **b,** The *J_comp_* values were compared between Clusters 1 and 2 in every cell type and visualized with a boxplot. Wilcoxon rank-sum tests were performed, and the Bonferroni method was applied for multiple comparison correction. In summary, STopover could classify multiple tissues into two distinct TME profiles. ns: not significant.

### STopover estimates spatial cell□cell interactions in barcode-based SRT of lung cancer

STopover can be applied to capture the overlapping location of ligand receptor (LR) expression. Based on the assumption that LR interaction mostly occurs between cells in proximity, the colocalized tissue domain extracted between LR pairs can be considered a key location for cell□cell interaction. The LR gene pairs provided by CellTalkDB^15^ and their expression profiles were utilized to estimate the key location and extent of the cell□cell interaction. Last, the strength of the interaction was ranked by *J_comp_* values between LR pairs. A representative tissue slide with high PD-L1 expression (*spa18ca02*) was selected, and the top LR pairs with *J_comp_* over 0.200 were extracted (**Supplementary Table 2**). The top 3 LR pairs with the highest ligand gene expression and *J_comp_* were *B2M-TFRC, B2M-KLRC1*, and *B2M-CD3G* (**Fig. 5a**). The main location for the LR pairs corresponded more to myeloid cells than to tS2 distribution (**Supplementary Fig. 6**). Next, an enrichment analysis was performed for the extracted LR pairs, and the top 10 Gene Ontology (GO) terms were extracellular matrix (ECM), MAPK/ERK pathway, cytokine-mediated signaling, and cell proliferation (**Fig. 5b**), which are closely related to cancer cell proliferation and metastasis^16^.

**Fig. 5.**
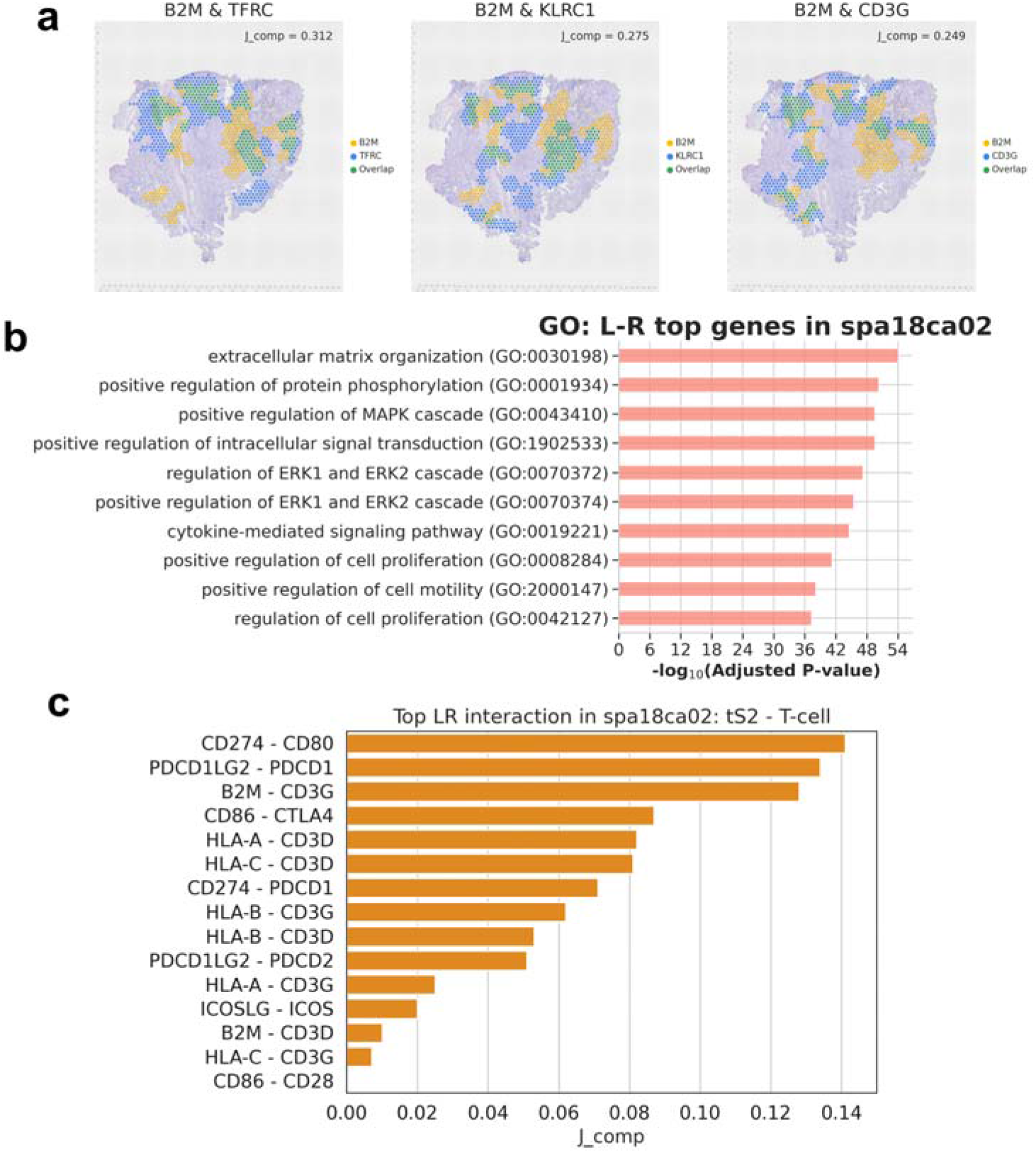
STopover predicts dominant cell–cell interactions in lung cancer tissue using barcode-based SRT. Based on the presumption that cell–cell communication mediated by LR interaction occurs in close proximity, spatial overlap patterns between the LR pairs were searched based on the CellTalkDB database^15^. The top LR pairs showing a high overlap score represented by *J_comp_* were considered dominant cell–cell interactions in the given tissue. The LR pairs with a *J_comp_* score over 0.2 were selected. **a,** Among the filtered LR pairs, the location of CCs for the top 3 pairs showing the highest average ligand gene expression in the tissue and the highest *J_comp_* value was mapped to the tissue. CC locations for features x and y are colored yellow and blue, respectively, and intersection locations are shown in green. **b,** Gene Ontology analysis was performed for all of the filtered LR pairs, and the enriched biological process terms are listed in ascending order of adjusted p values. To further investigate the cell–cell communication that occurs specifically between tS2 and T cells, 15 LR pairs closely related to T cell action were chosen, and their spatial colocalization patterns were extracted with STopover. Then, extracted CCs for LR pairs were intersected with the colocalized domain between tS2 and T cells, and the modified CCs were presumed to represent key locations for interaction between tS2 and T cells. **c**, *J_comp_* scores were calculated between the modified CCs, and the 15 LR pairs were listed in descending order of *J_comp_*. As a result, STopover could be adopted as a tool to screen dominant cell–cell interactions and their functional implications in cancer tissue.

To focus on the tumor-immune interaction in the selected high PD-L1 tissue, LR pairs related to T cell activation or suppression were chosen^17^. The CCs for the selected LR pairs were computed, and *J_comp_* was calculated for the spots corresponding to the tissue region where tS2 and T lymphocytes were colocalized. Among the LR pairs related to T cell action, *CD274-CD80* (*J_comp_*: 0.141), *PDCD1LG2-PDCD1* (*J_comp_*: 0.134), *B2M-CD3G* (*J_comp_*: 0.128), and *CD86-CTLA4* (*J_comp_*: 0.087) were the top gene pairs estimated to have high spatial overlap (**Fig. 5c and Supplementary Fig. 7**). This finding suggests the balance of T cell activation and inhibition signals at the T cell infiltrating tissue domain of the selected PD-L1 high tissue. In short, STopover can provide unbiased information on dominant spatial cell□cell interactions in lung cancer tissue and list the key components of cancer and T cell interactions.

### STopover extracts the spatial configuration of the lung cancer TME in image-based SRT

To prove the scalability of STopover in image-based SRT, the method was applied to the CosMx SMI platform in which the predefined RNA transcripts are detected by fluorescence signal, and cell-level RNA expression is calculated by cell segmentation. Lung cancer tissue was selected to investigate the TME. For convenience, the field of view (FOV) of CosMx data was divided into 100 by 100 grids, and RNA transcripts were assigned to each grid based on the 2D coordinates (**Supplementary Fig. 8a**). The fraction of a certain cell in a grid was determined by the total RNA counts belonging to the cell in each quadrant (**Supplementary Fig. 8b**). Then, by utilizing a cell-level RNA count matrix and a reference lung cancer single-cell dataset^13^, each cell could be classified into cell types defined from the single-cell data (**Supplementary Fig. 8c**). The abundance of a certain cell type was obtained by collecting the cells belonging to the cell type on each grid and calculating the sum of the cell fractions (**Supplementary Fig. 8d**). Additionally, cell-type specific expression was estimated by extracting the RNA transcript in each grid corresponding to the cell type. As a result, the image-based SRT was converted to grid-based expression data, and similar strategies were applied as with barcoding-based SRT to calculate overlapping spatial domains of feature pairs.

First, the spatial abundance of cell types (**Supplementary Fig. 9**) and spatial colocalization patterns between tS2 and major lung cancer cell types (**Fig. 6a**) were computed and visualized. Overall, the extracted CCs matched the spatial distribution pattern of cell types. When the global overlap score, *J_comp_*, was calculated, endothelial cells and fibroblasts ranked as the top 2 cell types, while *J_comp_* for immune cells was less than two-thirds of the fibroblasts (**Fig. 6a**). Notably, fibroblasts were mostly colocalized with tumors at the border regions. Additional STopover analysis was performed between tS2 and T cell subtypes (**Supplementary Fig. 10**). *J_comp_* scores were low across all cell subtypes, and CD8+ T cells, including naïve CD8+ T cells and cytotoxic CD8+ T cells, showed the lowest overlap with tS2 (*J_comp_*: 0.011 and 0.033).

**Fig. 6.**
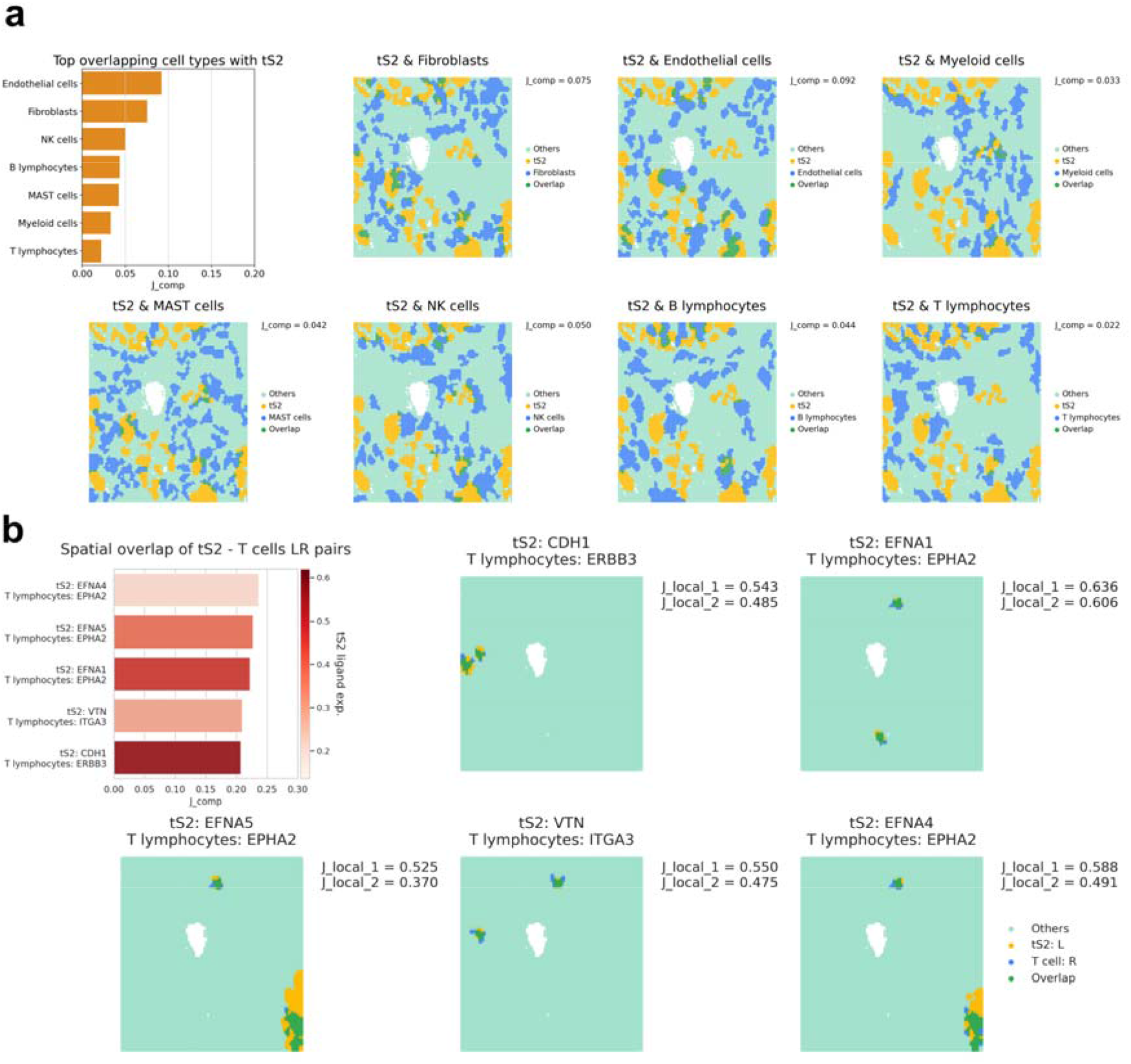
STopover captures cell colocalization patterns and estimates tumor and T cell crosstalk in lung cancer tissue using image-based SRT. The STopover was applied to image-based SRT of lung cancer tissue and deciphered spatial patterns of the cell types and their overlap. **a,** The *J_comp_* scores were calculated between tS2 and other main cell types and were visualized with a bar plot at the top left corner. The locations of CCs for tS2 (yellow) and other cell types (blue) were mapped to the tissue, and the overlapping domain is highlighted in green. The cell type-specific RNA counts were extracted from image-based SRT, and LR analyses were implemented using tS2-specific ligand expression and T cell-specific receptor expression. **b,** The *J_comp_* scores were computed for the LR pairs, and the results are presented in a bar plot in descending order of *J_comp_*. The average expression of ligand in tS2 was color-coded on the bar plot. Additionally, the location of the top 2 CC pairs showing the highest spatial overlap represented by *J_local_* was visualized on the grid. *J*_*local*_1_ is the highest *J_local_* score, and *J*_*local*_2_ is the second highest *J_local_* for the given LR pairs. In short, STopover can be flexibly applied to image-based SRT to calculate cell–cell colocalization patterns and predict key intercellular communications.

### STopover predicts T cell-specific cell□cell interactions in image-based SRT of lung cancer

STopover was applied to the lung cancer CosMx SMI dataset, and T cell-specific cell□cell interactions in the TME were explored. The list of LR pairs was refined by searching the intersection between genes present in the CosMx dataset and those from CellTalkDB. Then, ligand gene expression in tS2 cells and receptor gene expression in T cells were utilized to predict tumor-immune specific interactions. The analysis revealed 5 key interactions between tS2 and T cells with a *J_comp_* score over 0.200: *CDH1-ERBB3, EFNA1-EPHA2, EFNA5-EPHA2, VTN-ITGA3*, and *EFNA4-EPHA2* (**Fig. 6b and Supplementary Table 3**). Among the pairs, *CDH1-ERBB3* had the highest average ligand expression in the tissue. *CDH1* is a tumor suppressor and blocks EGFR signal transduction via cell contact^18^. The top 2 tissue subregions with the highest local overlap score, *J_local_*, were compared between the 5 LR pairs. Overall, the estimated location of the interaction between tS2 and T cells was highly overlapped across the selected LR pairs (**Fig. 6b**). In summary, STopover can be extensively used in image-based SRT to decipher the spatial patterns of the lung cancer microenvironment and rank the strong interaction specific to the tumor and T cells.

## Discussion

Decoding the spatial relationship between tumor, immune, and stromal cells in the TME is crucial to understanding the immune cell action on cancer cells and predicting responses to immunotherapy. Recent advances in SRT techniques have allowed for the screening of spatial patterns of multiple cell types and their gene expression in heterogeneous cancer tissue. In the case of barcoding-based spatial transcriptomics, the tissue is divided into small unit regions, and genome-wide RNA expression is investigated. This technique has been widely utilized because it can screen RNA expression across a wide range of tissues without predefined RNAs of interest^19–24^. Meanwhile, image-based spatial transcriptomics allows for spatial profiling of RNA expression in units of cells by identifying the RNA sequence, specifying the location using a fluorescence signal, and drawing the cell boundary^25–30^. If the two complementary spatial transcriptomic technologies are adopted, the spatial composition of the various cells constituting the tissue can be explored, and the complex interaction between the cells can be estimated.

In this regard, we developed STopover, which utilizes barcodes and image-based SRTs to summarize the topological colocalization pattern between cell types and LR pairs from CCs acquired by Morse filtration. By quantifying and visualizing the overlap between CC pairs, we compared the tumor infiltration of fibroblasts and immune cells in multiple cancer tissues. The key locations for cell infiltration were highlighted by ranking the local overlap of CCs between the feature pair. In addition, given that cell□cell interactions mediated by LR interactions occur in close proximity, major intercellular communication in cancer tissue could be estimated by finding the colocalization pattern of LR pairs. In particular, by utilizing cell type-specific RNA counts obtained from image-based SRT, STopover could list key spatial communication between tumor and T cells in lung cancer tissue.

One of the key features of STopover is capturing locally active regions of cell□cell colocalization in the TME. Compared to the thresholding-based approach to emphasize the overlapping tissue domain, STopover can readily spotlight the satellite regions where the abundance of cancer cells is relatively low, but immune cells are highly infiltrated (**Fig. 2**). Recently, a deep learning-based tissue image analyzer enabled the quantification of immune infiltration patterns in cancer tissues^31^. The main difference between this image analysis and STopover is that STopover provides an unbiased landscape of TME by exploring the infiltration pattern of all cell types present in the given tissue. Based on the functionality, STopover could dissect immune-excluded and immune-inflamed environments in one of the PD-L1 low and high tissues, respectively (**Fig. 3**). The TME of the two tissues was characterized by contrastive spatial overlap patterns of tumor-stromal and tumor-immune cells. The STopover was applicable not only in barcode-based SRT but also in image-based SRT platforms to understand the TME (**Fig. 6**). Moreover, the cross-cell type colocalization patterns in the tumor were represented by the global overlap score, *J_comp_*, and the lung cancer tissues could be classified into two clusters with distinct TME profiles (**Fig. 4**). The two clusters did not completely match the group divided by PD-L1 expression, implying that STopover can describe the configuration of the TME independently of PD-L1 levels in cancer. In addition, STopover offers biologically relevant information regarding immunotherapy response. As an example, the *J_comp_* score across the 11 tissues was positively correlated with the expression of genes coding MHC-class I protein (*B2M, HLA-A, HLA-B*,and *HLA-C*) (**Supplementary Fig. 5**), which are among the biomarkers for immunotherapy response^5^.

Another important functionality of STopover is that it can predict spatial tumor-immune interactions by ranking coexpression patterns of LR pairs in the whole tissue. One similar approach inferred cell□cell communication from the curated LR database by searching LR coexpression in the neighboring regions and measuring the number of distinct cell types^32^. Other approaches split the variability of gene expression into multiple factors ^33, 34^ or use graph neural networks^35, 36^ to model spatial cell□cell interactions. Compared to the suggested methods, STopover is a platform-agonistic method that can be utilized in both image- and barcode-based SRT and segment the key regions of the top-ranked cell□cell interaction. In one of the barcode-based SRT datasets, the top LR pairs with high spatial overlap were enriched with GO terms related to the development of the TME^16^ (**Fig. 5b**). In particular, the top 3 pairs were explained by cell□cell interactions via MHC class I molecules (**Fig. 5a**), and their main location of interaction was highly overlapped with the domain of myeloid cells but exclusive to tS2 (**Supplementary Fig. 6**). This finding implies that the MHC class I-mediated process, which is one of the most activated cell□cell communications, occurs primarily at the myeloid cell niche. In the case of the image-based SRT dataset, tumor- and T cell-specific expression profiles could be extracted, and the tumor and T cell interaction could be more specifically investigated (**Fig. 6b**). A total of 5 LR pairs were selected as meaningful interactions, and there was not enough literature evidence that highlights the selected interaction in the TME. This is explainable considering that the given cancer tissue has a low immune infiltration and that the extracted interaction could be a minor portion of the whole tissue. The main niche for tumor and T cell communication was highly redundant and constrained within small regions (**Fig. 6b**), further explaining the T cell exclusive TME.

There are several points to consider when applying STopover to dissect the TME. First, caution is needed when interpreting the top cell□cell interaction extracted by STopover in barcode-based SRT. Because the LR interaction is searched for all the cell types present in the unit domain of the tissue, the highlighted location does not indicate the specific interaction between the two cell types. An alternative approach to estimate cell type-specific interactions might be constraining the CCs of LR pairs to the spatial overlapping niche between the two cell types of interest. Because the cell type-specific expression can be extracted in image-based SRT, the suggested approach could be validated. Compared to the key location of tS2 and the T cell-specific interaction of the top LR pairs (**Supplementary Fig. 11a**), the regions extracted by the alternative approach were similar (**Supplementary Fig. 11b**). Last, there may be a concern that the range of cell□cell interactions is not considered and that the LR colocalization pattern extracted from STopover cannot differentiate short-range from long-range cell□cell communication. However, because the diameter of the spot, the basic unit of the Visium dataset, is 55 micrometers and the size of the generated grid in the CosMx dataset is 49.214 by 39.376 micrometers, long-range interactions in the CellTalkDB database, such as cytokine and receptor signaling (characteristic length scale: ~100 micrometers)^37^, were presumed not to exceed the range of a few spots or grids.

In summary, STopover is a robust tool that utilizes SRT and topological analysis to analyze the spatial infiltration patterns of the TME and highlights the key niche of tumor-immune and tumor-stromal interactions. The proposed tool is expected to be applicable to elucidate immune evasion mechanisms and guide patient-specific strategies to enhance the efficacy of ICI treatment in cancer.

## Methods

### Simulation dataset

The simulation dataset was created to test the usefulness of STopover in capturing locally active subregions where one of the features has a low value while the other has a high value. For simplification, the spatial map of two features, tumor and immune cells, was created by adding a trimmed 2D Gaussian function multiple times to the 100 by 100 grid with zero background.

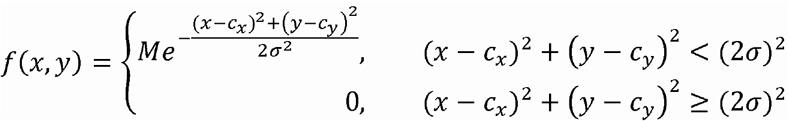

*f(x,y): trimmed 2D Gaussian function for the simulated feature value, M: maximum feature value in the center, c_x_: x coordinate of the center, c_y_: y coordinate of the center, σ: standard deviation*

The input values, M, c_x_, c_y_, and σ, were (8, 30, 30, 20), (8, 60, 70, 10), (2, 70, 20, 10), and (1, 90, 90, 10) in tumor cells and (4, 95, 85, 5), (2, 20, 20, 5), (2,70, 20, 5), (1, 70, 70, 5), and (1, 85, 95, 5) in immune cells. Then, the noise was generated by random sampling from a Gaussian distribution (mean: 1 and standard deviation: 0) and multiplying the sampled values by 0.01. Last, to mimic the noise in the SRT dataset, two independently generated noise samples were added to the 2D grid of tumor and immune cells, and the negative values were replaced with zero.

### Barcode-based SRT of human lung cancer

Eleven lung adenocarcinoma tissue samples were acquired from 7 patients who were initially diagnosed with lung cancer and underwent surgical resection. The patient number and the tissue number were used to name the samples. For example, the second tissue obtained from patient number 18 was named spa18ca02. The samples were divided into two groups based on the PD-L1 expression level. The PD-L1 low group was composed of spa01ca01, spa01ca02, spa02ca01, spa02ca02, spa06ca01, and spa10ca01 with PD-L1 expression of 0. The PD-L1 high group expression group was composed of spa16ca01, spa17ca01, spa17ca02, spa18ca01, and spa18ca02 with PD-L1 expression above 80%. The acquired samples were embedded in an optimal cutting temperature (OCT) compound, cryosectioned, and processed to generate Visium spatial transcriptomic datasets. The diameter of the spot, the basic unit of the Visium spatial transcriptome, was 55 micrometers, and the distance between the spots was 110 micrometers. The number of spots and genes utilized for the analysis, patient history, and histological information of the samples are summarized in **Supplementary Table 1**. The cell type composition in the spot was inferred based on the reference single-cell data obtained from human lung cancer tissue^13^. The cell types were classified into 8 large categories (epithelial cells, fibroblasts, endothelial cells, myeloid cells, MAST cells, NK cells, B lymphocytes, and T lymphocytes) and 47 cell subtypes. For example, the T lymphocytes were divided into naïve CD4+T, CD4+ Th, CD8+/CD4+ mixed Th, exhausted Tfh, Treg, naïve CD8+ T, cytotoxic CD8+ T, exhausted CD8+ T, CD8 low T cells, and T lymphocytes_ns.

### Image-based SRT of human lung cancer

The publicly available CosMx SMI dataset^38^, which is one of the image-based SRT platforms, was utilized for STopover analysis. The dataset was a formalin-fixed paraffin-embedded (FFPE) sample of non-small cell lung cancer. The RNA was captured across 30 FOVs, and the size of each FOV was 0.985 by 0.657 millimeters. The margin of the cells was segmented based on the immunofluorescence image, and the cell-level count matrix was constructed. The coordinate of the transcript, cell assignment data of each transcript, cell-level expression profiles, and cell metadata files were utilized for the STopover analysis. The numbers of transcripts and cells were 30,370,769 and 100,149, respectively. Among the transcripts, 37,226,610 could be assigned to the cell and corresponded to 960 gene symbols and 20 negative probes, which do not correspond with any sequence. The cell type annotation of the segmented cells was performed with identical lung cancer single-cell data^13^, which were used for cell type decomposition in barcode-based SRT.

### Preprocessing barcode- and image-based SRT

The preprocessing steps for the SRT datasets are mainly based on Scanpy (ver. 1.9.1)^39^ running on Python (ver. 3.8). First, the spot-level count matrix of barcode-based SRT was processed for downstream analysis. The RNA count of each spot was normalized such that the total count became 10,000 and log-transformed [ln (normalized count +1)]. Cell type decomposition was performed by applying the CellDART algorithm^14^ to each Visium spatial transcriptomic dataset based on the reference single-cell dataset. As a result, a spatial composition map of all cell types in the lung cancer tissue was generated. The spatial distribution of cell types was utilized to calculate cell□cell colocalization patterns. Additionally, the spatial expression of LR pairs was adopted to estimate spatial cell□cell communication.

Second, image-based SRT data were processed to create a grid-based count matrix, which enabled analysis of the data similar to the barcode-based SRT. The whole FOV, including all 30 small FOVs, was divided into 100 by 100 grids based on the outermost coordinate of the transcript (**Supplementary Fig. 8a**). The size of the unit grid is approximately 49.214 by 39.376 micrometers, which is similar to the diameter of the spot from Visium spatial transcriptomics. The grid-level RNA count was normalized to fit the total count in each cell to 1,000 and log-transformed. The fraction of the cell assigned to each grid was defined by the ratio of total RNA counts in a portion of the cell belonging to the grid to total RNA counts in the cell (**Supplementary Fig. 8b**). Then, the cell-level count matrix was utilized to annotate the cells based on the reference single-cell data with the ‘Ingest’ algorithm provided by Scanpy (scanpy.tl.ingest)^39^ (**Supplementary Fig. 8c**). The tool maps the reference single-cell expression data into the spatial single-cell embedding using principal component analysis (PCA) and the k-nearest neighbor (kNN) search method suggested in the uniform manifold approximation and embedding (UMAP) algorithm^40^. After the annotation of cell types, the grid-level abundance of cell types was calculated by summing the fraction of all cells in the grid corresponding to the cell type (**Supplementary Fig. 8d**). Of note, cell type-specific expression was calculated by extracting the transcripts belonging to the specific cell types and performing log-transformation of the summed count. Finally, a spatial map of cell type abundance was utilized to calculate spatial cell□cell localization and grid-level log-normalized counts to compute spatial cell□cell interactions.

### STopover: extracting colocalized patterns of a feature pair

STopover applies Morse filtration and the dendrogram smoothing algorithm^11^, one of the topological analysis methods, to extract CCs from the given feature pair and calculate the overlap between the CC pairs (**Fig. 1 and Supplementary Fig. 1**). First, to reduce the intrinsic sparsity of features in SRT, a Gaussian smoothing filter was applied to the spatial feature map. The filter with a full-width half maximum (FWHM) of 2.5 times the unit central distance between spots or grids was applied for smoothing. Next, CCs were calculated for all thresholds while lowering the value from the highest feature value in the tissue to the lowest value using the algorithm suggested by NetworkX^41^. The existing CCs from the higher threshold were gradually merged or new CCs were defined based on the spatial distance from the newly added spot or grid to the existing CCs. This hierarchical clustering process was summarized with a dendrogram. Each vertical bar in the dendrogram represents the start and end of threshold values that a certain CC is continuously observed, and each horizontal bar links the existing CCs (child) with merged or new CCs (parent) from the lower threshold value. Then, the dendrogram was smoothed such that the hierarchical structure of the spatial feature map could be simplified. The vertical bars are selected in the order of the longest length to the shortest length, and if the connected children CCs of the selected bar have a smaller size than the minimum size of CCs, then their elements are removed and aggregated to the parent. Additionally, the start point of the vertical bars is updated to have the maximum feature value in the CC. The process is iterated until all of the vertical bars are selected and the uppermost bars of the dendrogram, reconfigured CCs, represent locally activated regions of the given feature. Next, the average feature value was computed within each CC region, and CCs, which are presumed to capture noise due to a low average value below a certain percentile, were deleted. Then, the Jaccard index was calculated for all possible pairs of reconfigured CCs from the two features and named *J_local_*. The CC pair with a high *J_local_* value indicates the tissue subregion where the extent of overlap between two features is high. Additionally, all CCs of each feature were aggregated, and the Jaccard index between the two aggregated CCs was calculated and named *J_comp_*. The formulas below explain how the Jaccard indices were calculated.

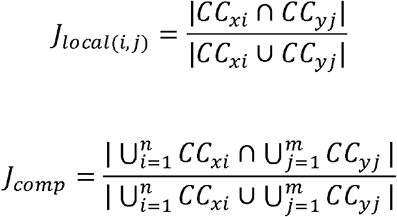

*CC_xi_: set including spots or grids belonging to the i-th connected component of feature x, CC_yj_: set including spots or grids belonging to the j-th connected component of feature y*

The main parameters in the models are the minimum size of CCs (s_min_) expressed as the number of spots or grids and the percentile threshold to remove the CCs with low average feature values (p_t_). Increasing both s_min_ and p_t_ is expected to remove the noise. However, if s_min_ is too large, then too many CCs are aggregated, and if p_t_ is too large, then many CCs unrelated to noise will disappear, which impairs the performance of the STopover. The CCs and the overlap patterns between tS2 (tumor subtype) and T cells were examined across different s_min_ and p_t_ values (**Supplementary Fig. 12**). The optimal value of s_min_ was fixed to 20, considering that it corresponds to diameters of approximately 479.264 and 295.284 micrometers in circular CCs of barcode- and image-based SRTs, respectively, which is suitable for investigating subregional transcriptomic changes. Additionally, an optimal value of p_t_ depends on the noise level of the dataset, and by setting the FWHM of the Gaussian smoothing filter to 2.5, a p_t_ value of 30 was adequate to reflect the spatial patterns of the feature and remove noise (**Supplementary Fig. 12**). Because the spot in the barcode-based SRT and grid in the image-based SRT are similar in size, the optimal values of s_min_ and p_t_ in both datasets could be shared. In the case of the simulation dataset, the optimal s_min_ and p_t_ values to reduce the noise and better represent the spatial feature map were 20 and 80, respectively (**Supplementary Fig. 13**).

### STopover: estimating cell□cell interaction patterns in tumors

Cell□cell communication was estimated based on the assumption that cell□cell interactions mediated by LR interactions occur within a range of few spots or grids. CellTalkDB, the curated LR database, was selected^15^, and the spatial colocalization pattern of all LR expressions was searched. The LR pairs with a high colocalization score (*J_comp_*) were presumed to show high interaction in the tissue subregions represented by CCs. The most meaningful LRs were selected by removing the *J_comp_* below the threshold of 0.200. GO analysis^42, 43^ was performed for the filtered ligand and receptor gene sets using gseapy (version 0.10.8)^44, 45^. The overrepresented biological process terms were extracted to comprehend the functional role of the dominant LR interaction in the given tissue.

To search cell type-specific LR interactions in barcode-based SRT, modified CCs were defined as intersecting subregions between CCs obtained from LR interaction analysis and the colocalized tissue domain of the two cell types extracted by STopover. *J_local_* and *J_comp_* were calculated between the modified CCs of the two features. In the case of image-based SRT, the cell type-specific (cell types A and B) log-normalized count was first computed, and spatial overlap patterns between ligand expression of cell type A and receptor expression of cell type B were extracted to predict spatial cell□cell interactions.

## Code availability

Python source code and application for STopover were uploaded on https://github.com/bsungwoo/STopover.

