## Supplementary Fig for "STopover captures spatial colocalization and interaction in the tumor microenvironment using topological analysis in spatial transcriptomics data"

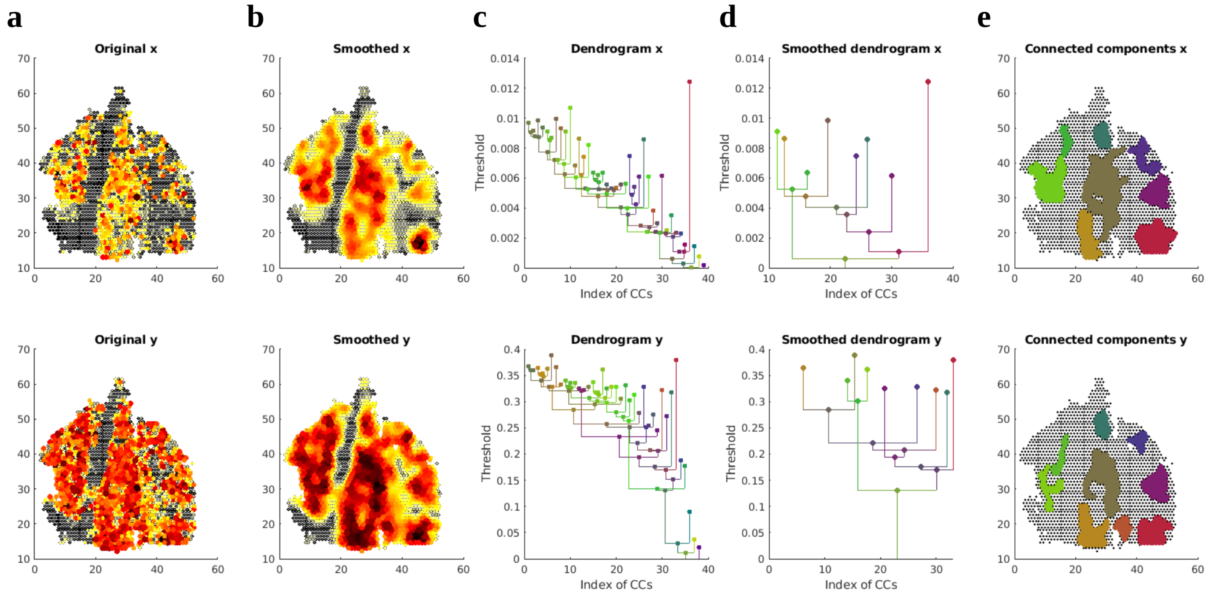


**Supplementary Fig. 1. Extracting CCs by Morse filtration and dendrogram smoothing in Visium dataset.**

An example of extracting CCs from a spatial map of feature x (tS2) and feature y (T lymphocytes), which is the core algorithm of STopover. The utilized Visium dataset is one of the PD-L1 high tissues (spa18ca02). (a, b) First, the original spatial maps of features were smoothed by applying a Gaussian filter with a full-width half maximum (FWHM) of 2.5 times the unit distance between the center of the spots. (c) Then, the CCs of both features were discovered based on the spatial distance between spots by gradually lowering the feature values from the highest possible threshold value to the lowest value. (d) The resulting CCs expressed as dendrograms were smoothed based on the minimum size of the CCs and the persistence of each CC during the threshold lowering. (e) Finally, reconfigured CCs, the uppermost bars of the dendrograms, were considered locally activated regions of the feature x and y.

To remove the noise of the spatial map, the CCs with the feature value below the certain percentile threshold were removed and the local and composite Jaccard indexes were calculated between the final CCs of the feature pair (feature x and y).

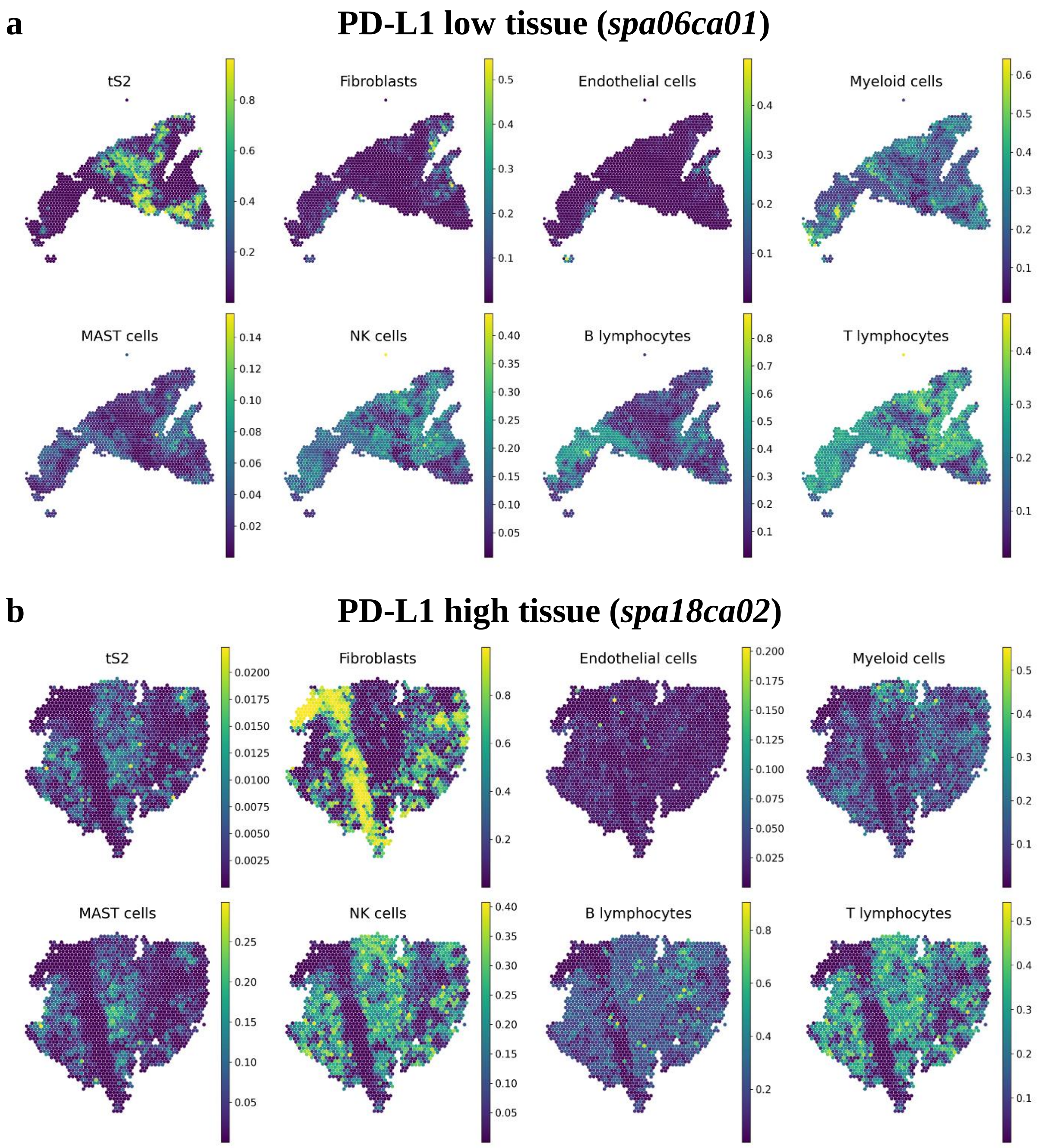
a
PD-L1 low tissue (*spa06ca01*)
tS2
Fibroblasts
Endothelial cells
Myeloid cells
MAST cells
NK cells
B lymphocytes
T lymphocytes
b
PD-L1 high tissue (*spa18ca02*)
tS2
Fibroblasts
Endothelial cells
Myeloid cells
MAST cells
NK cells
B lymphocytes
T lymphocytes

**Supplementary Fig. 2: Spatial distribution of tS2 and other main cell types composing the lung cancer tissue in barcode-based SRT.**

The spatial composition of tS2 and other main cell types (fibroblasts, endothelial cells, myeloid cells, MAST cells, NK cells, B lymphocytes, and T lymphocytes) in the representative (a) PD-L1 low (spa6ca01) and (b) high tissues (spa18ca02) were estimated and visualized.

**a**

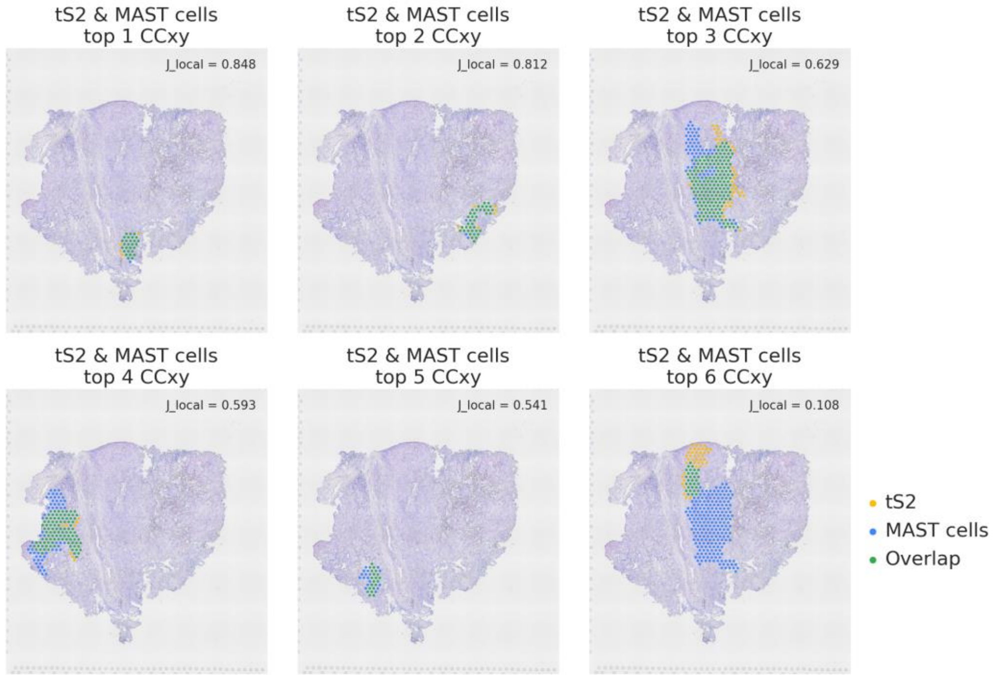
tS2 & MAST cells
top 1 CCxy
J_local = 0.848
tS2 & MAST cells
top 2 CCxy
J_local = 0.812
tS2 & MAST cells
top 3 CCxy
J_local = 0.629
tS2 & MAST cells
top 4 CCxy
J_local = 0.593
tS2 & MAST cells
top 5 CCxy
J_local = 0.541
tS2 & MAST cells
top 6 CCxy
J_local = 0.108
tS2
MAST cells
Overlap

**b**

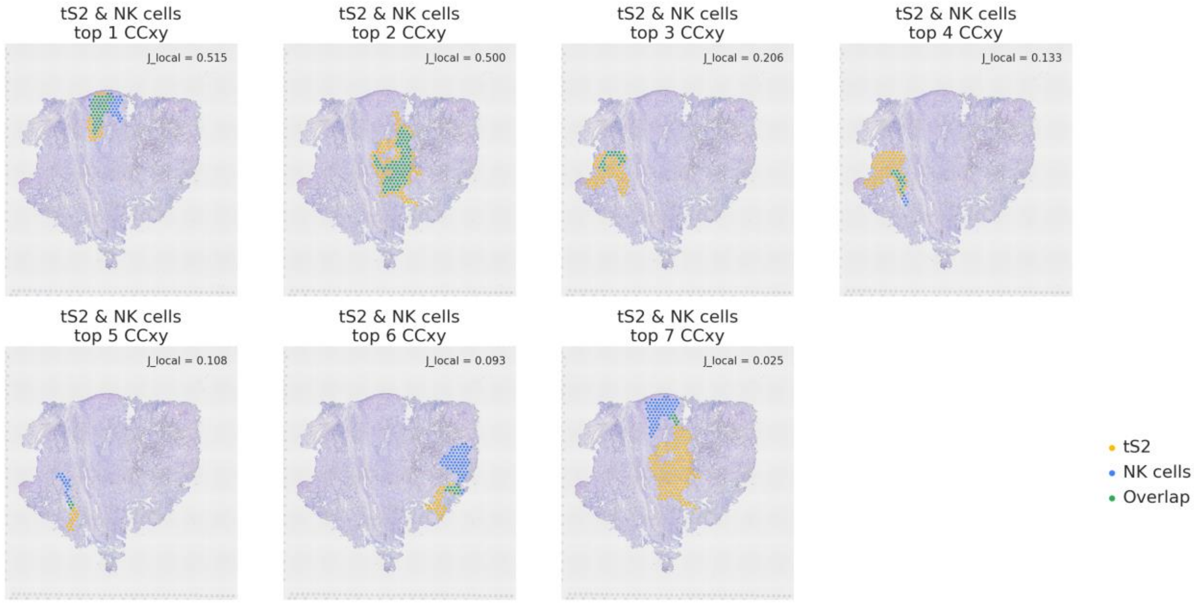
tS2 & NK cells
top 1 CCxy
J_local = 0.515
tS2 & NK cells
top 2 CCxy
J_local = 0.500
tS2 & NK cells
top 3 CCxy
J_local = 0.206
tS2 & NK cells
top 4 CCxy
J_local = 0.133
tS2 & NK cells
top 5 CCxy
J_local = 0.108
tS2 & NK cells
top 6 CCxy
J_local = 0.093
tS2 & NK cells
top 7 CCxy
J_local = 0.025
tS2
NK cells
Overlap

c

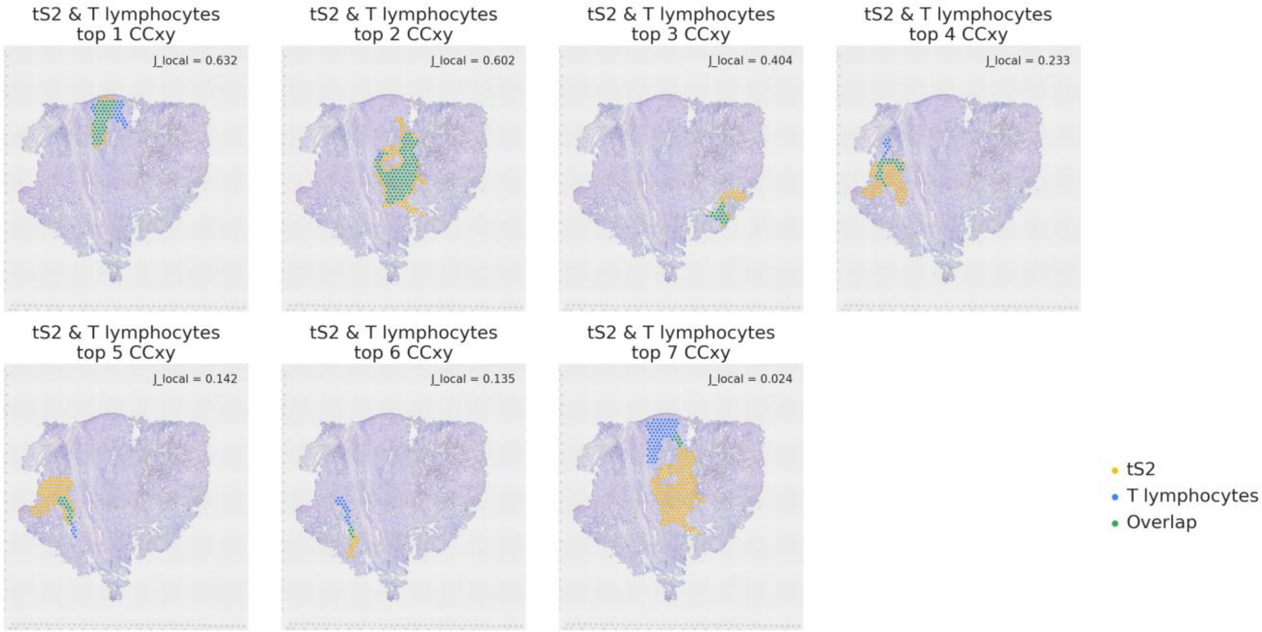


**Supplementary Fig. 3: The location of top CC pairs between tS2 and T lymphocytes in PD-L1 high lung cancer tissue (*spa18ca02*)**

The CC pairs between tS2 and (a) MAST cells, (b) NK cells, or (c) T lymphocytes were extracted by STopover and the top 7 pairs were visualized in the descending order of $J_{local}$ values.

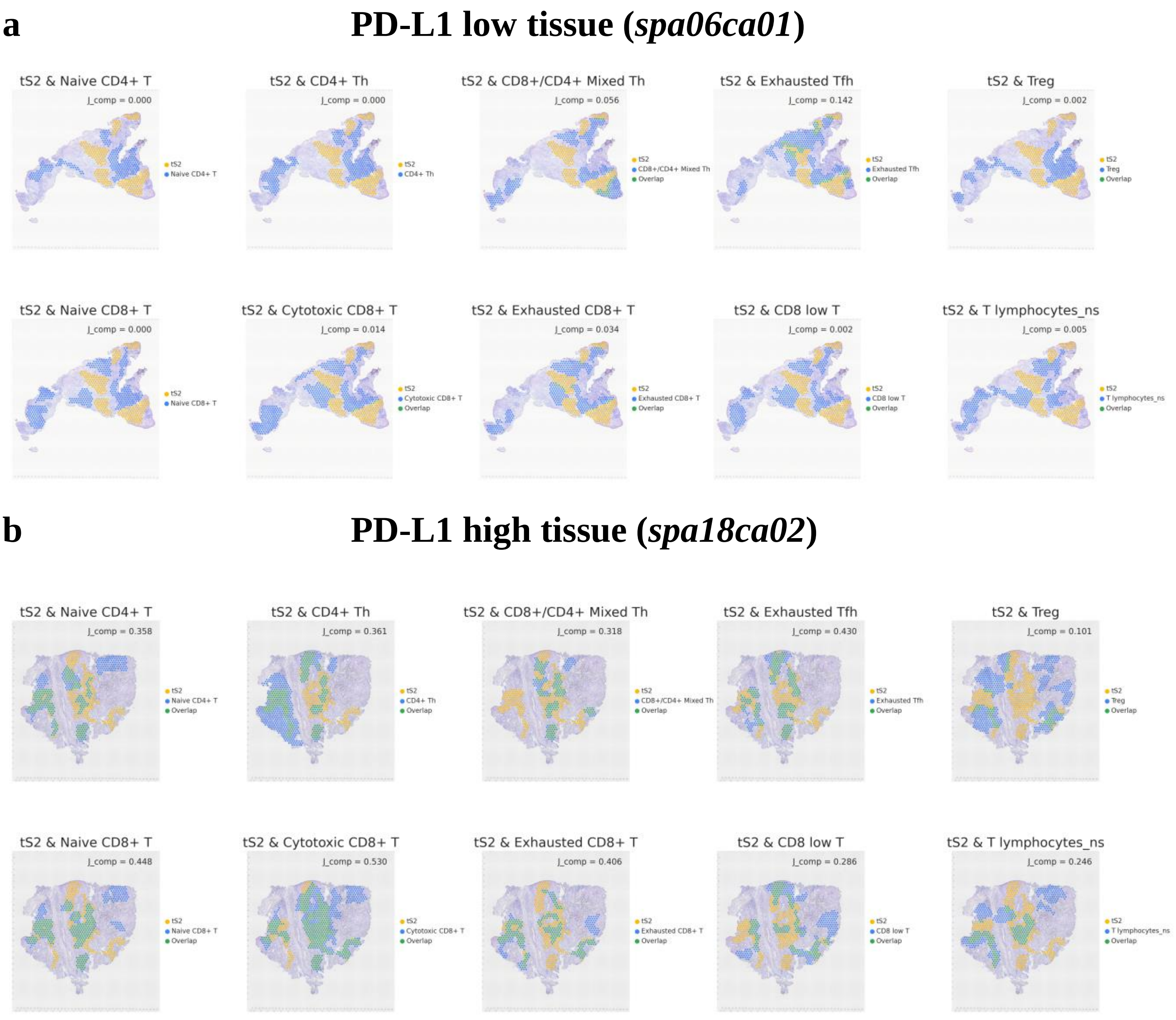


**Supplementary Fig. 4: The colocalization patterns between tS2 and multiple T cell subtypes in barcode-based SRT of lung cancer**

The key location of tS2 and multiple T cells subtypes were represented by CCs and the overlapping tissue domains were highlighted as the intersecting subregions between the two aggregated CCs. The analysis was performed in the two representative (a) PD-L1 low (*spa6ca01*) and (b) high tissues (*spa18ca02*).

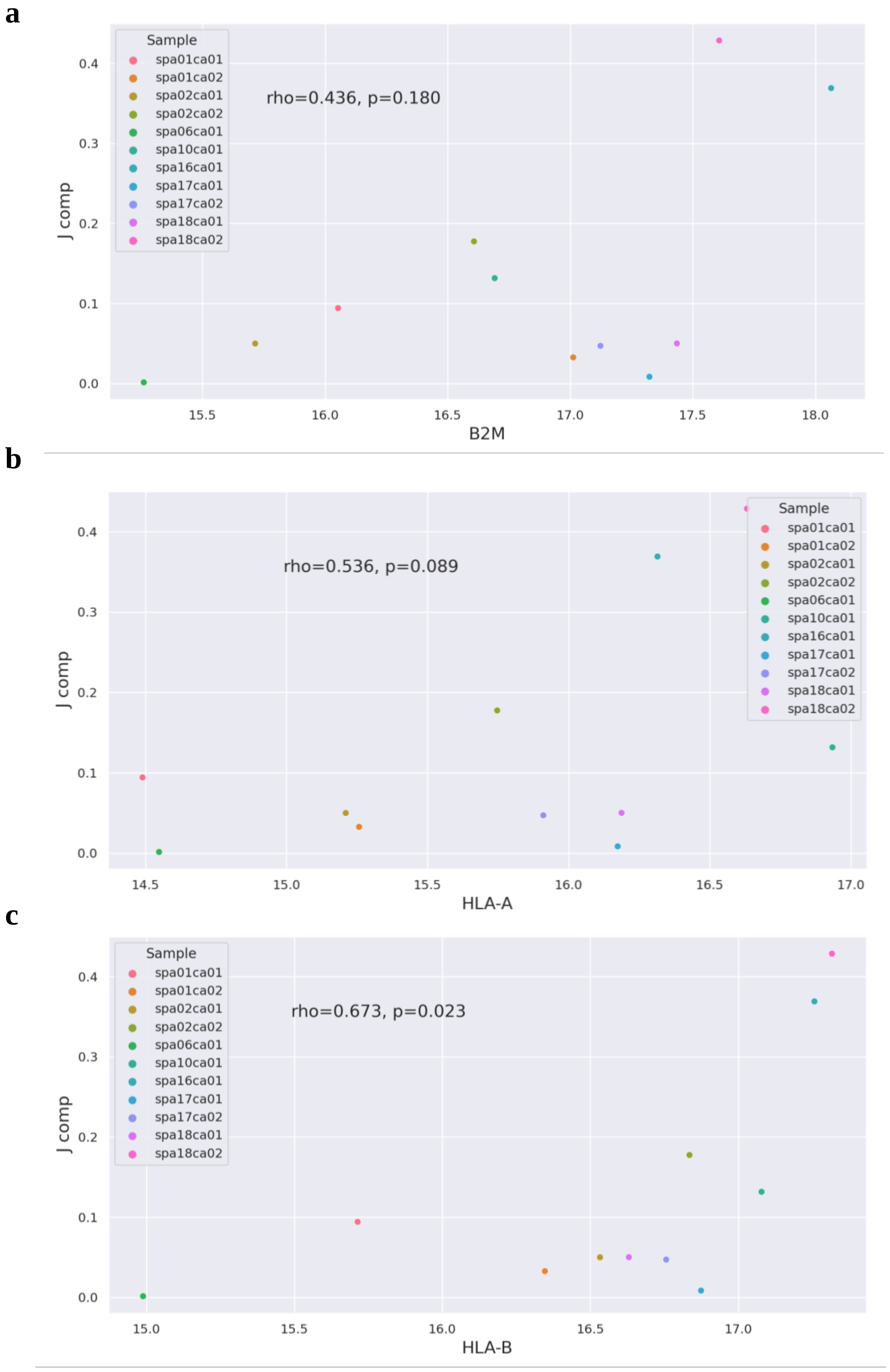
a
Sample
spa01ca01
spa01ca02
spa02ca01
spa02ca02
spa06ca01
spa10ca01
spa16ca01
spa17ca01
spa17ca02
spa18ca01
spa18ca02
rho=0.436, p=0.180
J comp
0.0
0.1
0.2
0.3
0.4
15.5
16.0
16.5
17.0
17.5
18.0
B2M
b
Sample
spa01ca01
spa01ca02
spa02ca01
spa02ca02
spa06ca01
spa10ca01
spa16ca01
spa17ca01
spa17ca02
spa18ca01
spa18ca02
rho=0.536, p=0.089
J comp
0.0
0.1
0.2
0.3
0.4
14.5
15.0
15.5
16.0
16.5
17.0
HLA-A
c
Sample
spa01ca01
spa01ca02
spa02ca01
spa02ca02
spa06ca01
spa10ca01
spa16ca01
spa17ca01
spa17ca02
spa18ca01
spa18ca02
rho=0.673, p=0.023
J comp
0.0
0.1
0.2
0.3
0.4
15.0
15.5
16.0
16.5
17.0
HLA-B

d

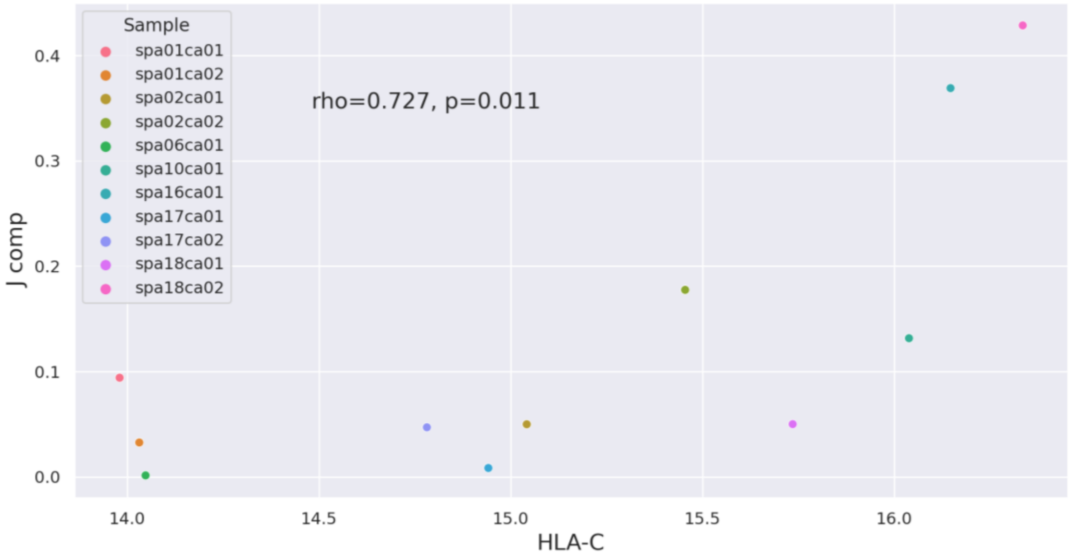

rho=0.727, p=0.011
Sample
spa01ca01
spa01ca02
spa02ca01
spa02ca02
spa06ca01
spa10ca01
spa16ca01
spa17ca01
spa17ca02
spa18ca01
spa18ca02
J comp
HLA-C


e

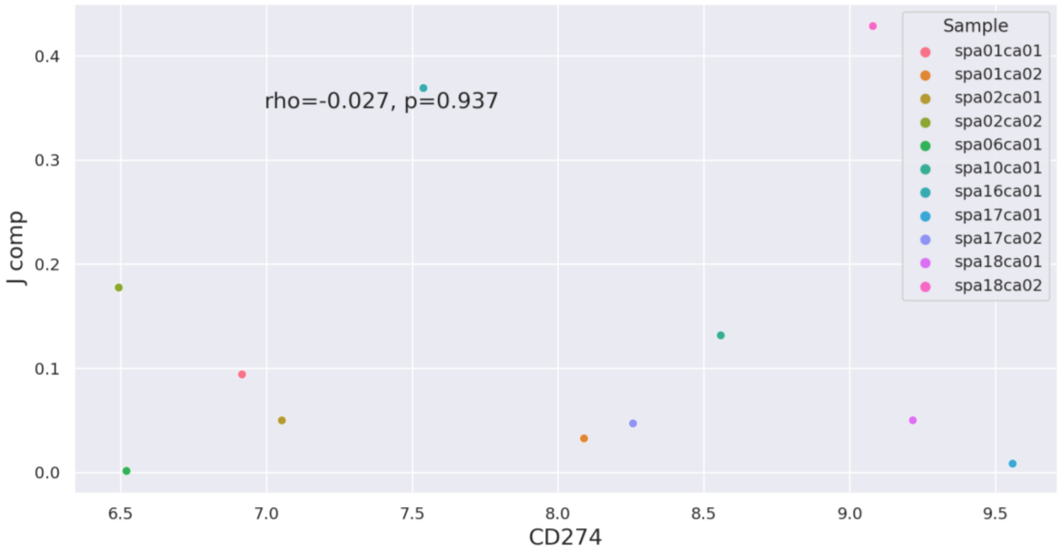

rho=-0.027, p=0.937
Sample
spa01ca01
spa01ca02
spa02ca01
spa02ca02
spa06ca01
spa10ca01
spa16ca01
spa17ca01
spa17ca02
spa18ca01
spa18ca02
J comp
CD274


f

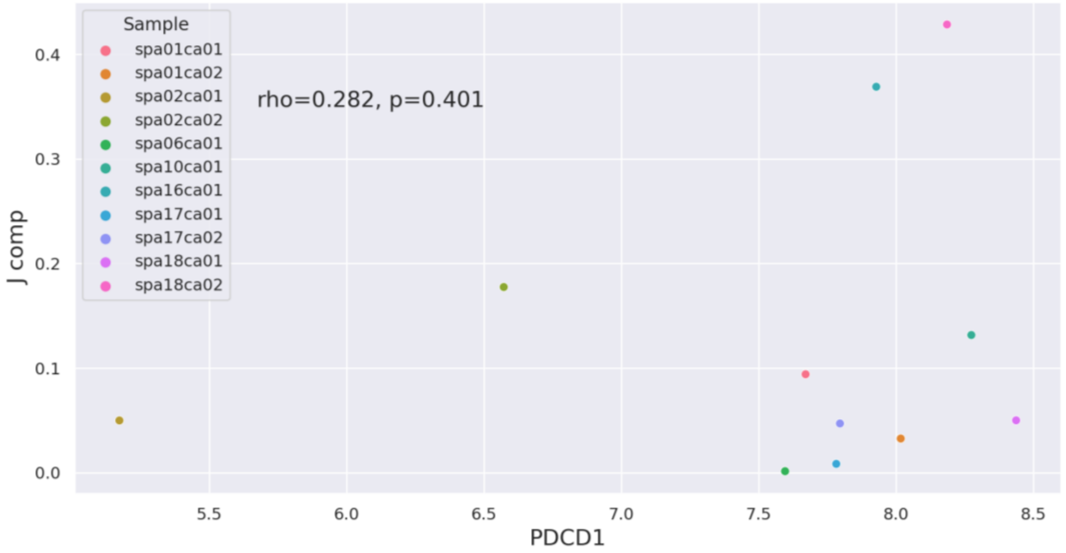

rho=0.282, p=0.401
Sample
spa01ca01
spa01ca02
spa02ca01
spa02ca02
spa06ca01
spa10ca01
spa16ca01
spa17ca01
spa17ca02
spa18ca01
spa18ca02
J comp
PDCD1

**Supplementary Fig. 5. Scatter plot between pseudobulk RNA expression and $J_{comp}$ value of tS2 and T cells**

The scatter plot shows the correlation between the pseudobulk RNA expression of (a) B2M, (b) HLA-A, (c) HLA-B, (d) HLA-C, (e) CD274, and (f) PDCD1, which was calculated by log-transforming total RNA counts, and the $J_{comp}$ value which represents the extent of spatial overlap between tS2 and T cells. The plot visualizes the data points obtained from 11 lung cancer tissues. The Spearman's correlation coefficient was computed and shown in the top right corner of each plot.

**a**

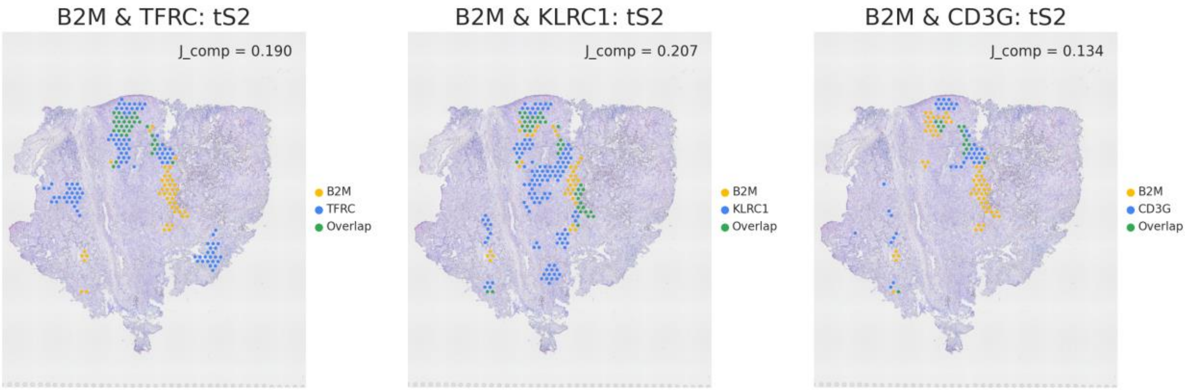


**b**

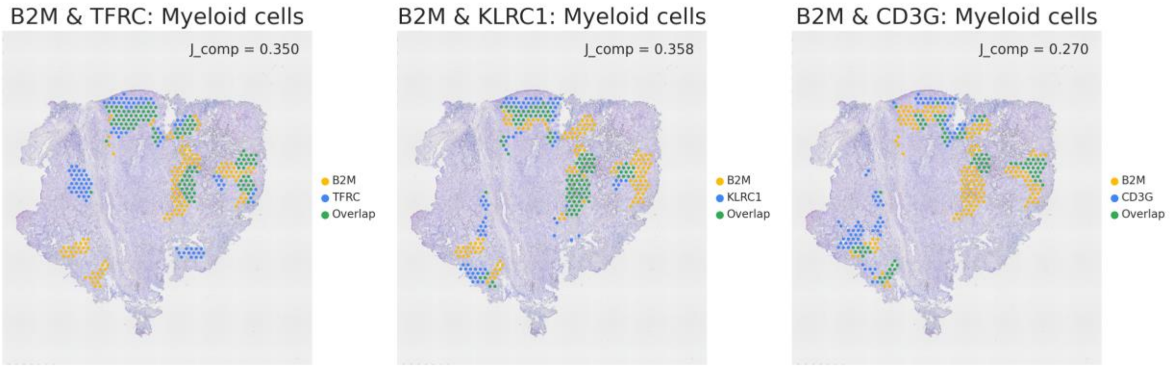


**Supplementary Fig. 6. The spatial overlap patterns of the top LR pairs constrained to the tS2 or myeloid cell domain in PD-L1 high lung cancer tissue (*spa18ca02*)**

The aggregated CCs were extracted for the top 3 LR pairs (*Jcomp* > 0.2) showing the highest ligand gene expression in the tissue and the highest $J_{comp}$ value. Then, the intersecting regions between the two aggregated CCs and CCs calculated from (a) tS2 or (b) myeloid cells were extracted. The location of modified CCs was mapped to the tissue and *Jcomp* values were computed with the two modified CCs.

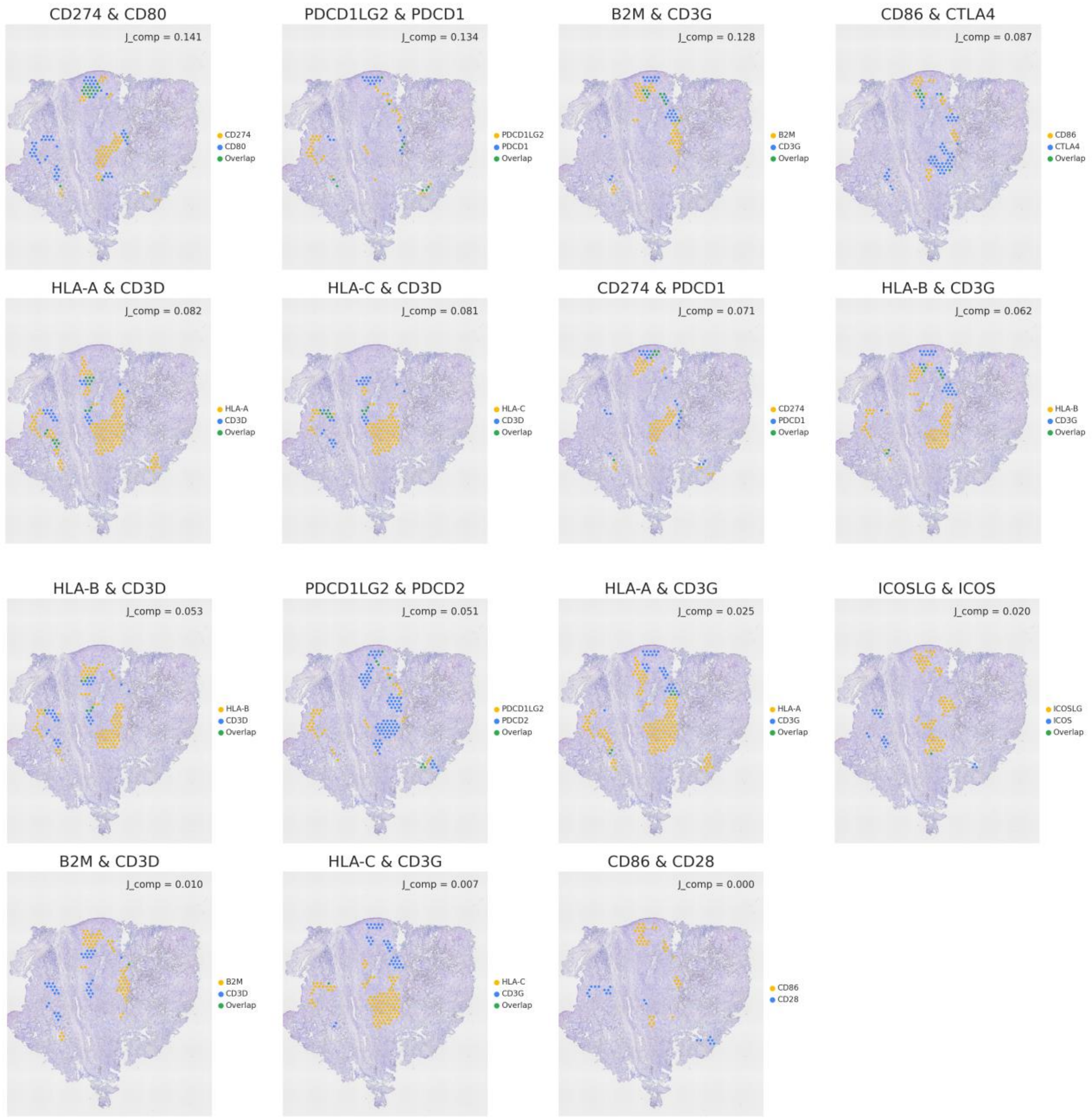
CD274 & CD80
J_comp = 0.141
CD274
CD80
Overlap
PDCD1LG2 & PDCD1
J_comp = 0.134
PDCD1LG2
PDCD1
Overlap
B2M & CD3G
J_comp = 0.128
B2M
CD3G
Overlap
CD86 & CTLA4
J_comp = 0.087
CD86
CTLA4
Overlap
HLA-A & CD3D
J_comp = 0.082
HLA-A
CD3D
Overlap
HLA-C & CD3D
J_comp = 0.081
HLA-C
CD3D
Overlap
CD274 & PDCD1
J_comp = 0.071
CD274
PDCD1
Overlap
HLA-B & CD3G
J_comp = 0.062
HLA-B
CD3G
Overlap
HLA-B & CD3D
J_comp = 0.053
HLA-B
CD3D
Overlap
PDCD1LG2 & PDCD2
J_comp = 0.051
PDCD1LG2
PDCD2
Overlap
HLA-A & CD3G
J_comp = 0.025
HLA-A
CD3G
Overlap
ICOSLG & ICOS
J_comp = 0.020
ICOSLG
ICOS
Overlap
B2M & CD3D
J_comp = 0.010
B2M
CD3D
Overlap
HLA-C & CD3G
J_comp = 0.007
HLA-C
CD3G
Overlap
CD86 & CD28
J_comp = 0.000
CD86
CD28

**Supplementary Fig. 7. The spatial overlap patterns of the T cell action-related LR pairs constrained to the tS2 and T cell colocalized domain in PD-L1 high lung cancer tissue (*spa18ca02*)**

The aggregated CCs were extracted for the T cell action-related LR pairs. Then, the intersecting regions between the two aggregated CCs and colocalized domain of tS2 and T cells were extracted. The location of modified CCs was mapped to the tissue and $J_{comp}$ values were computed with the two modified CCs.

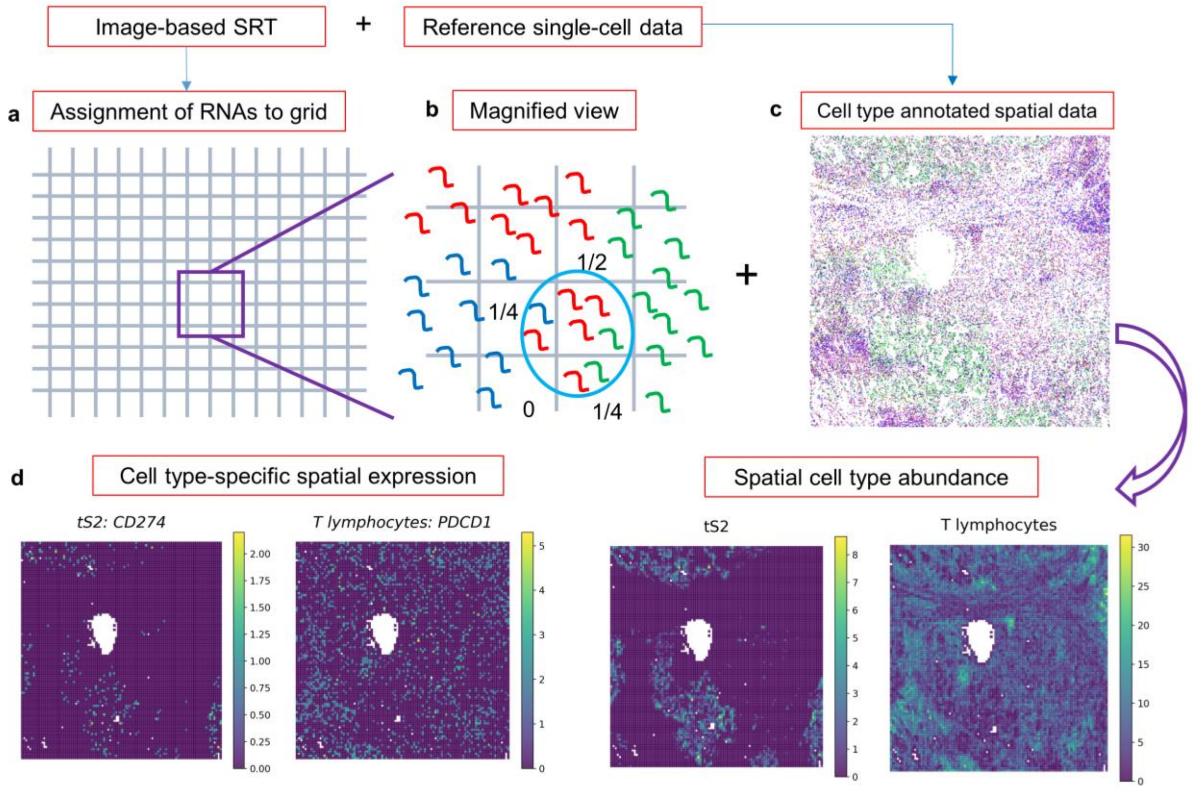


**Supplementary Fig. 8. Schematic image for converting image-based SRT to grid-based data**

(a) A whole field of view of image-based SRT was divided into grids (in the case of the CosMx SMI platform, 100 by 100 grids are applied) such that the spatial resolution can be adjusted similarly to the Visium spatial transcriptomic dataset. (b) Then, the RNA transcript was assigned to the grid and the fraction of a certain cell in a grid was calculated based on the total number of RNAs belonging to a certain quadrant of the grid. (c) Next, reference single-cell RNA-seq data was utilized to annotate the cells segmented in the image-based SRT. (d) By summing all the fractions of cells corresponding to a certain cell type, a spatial map of cell type abundance could be acquired. Also, cell type-specific spatial expression could be calculated by extracting RNAs belonging to a certain cell type.

**a**

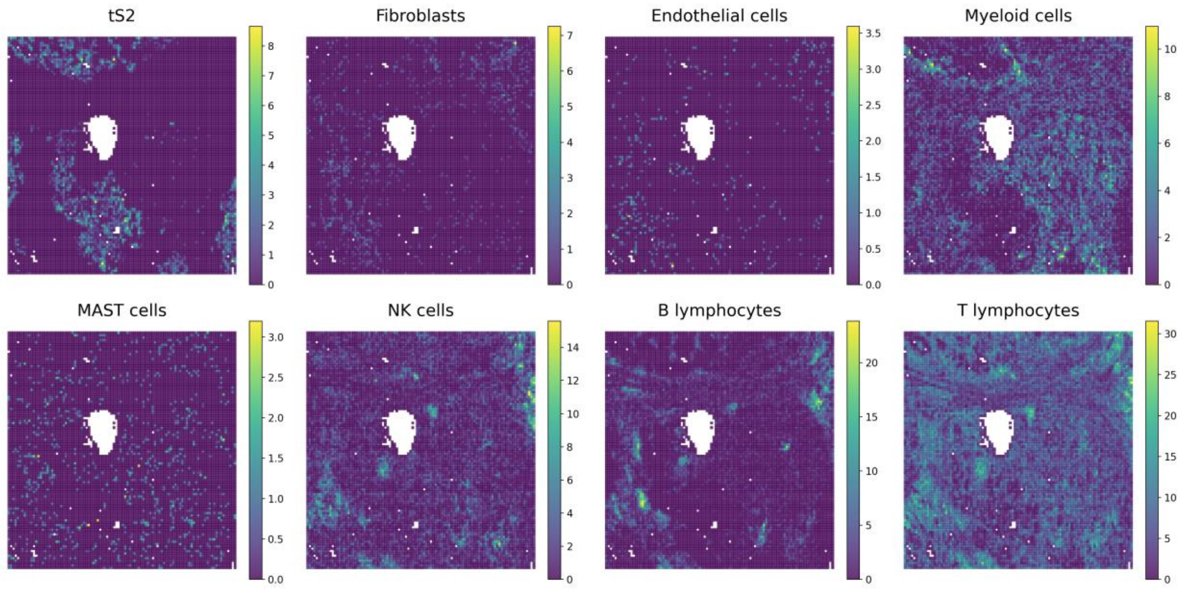


**Supplementary Fig. 9. Spatial distribution of tS2 and other main cell types composing the lung cancer tissue in image-based SRT.**

The spatial map for an abundance of tS2 and other main cell types (fibroblasts, endothelial cells, myeloid cells, MAST cells, NK cells, B lymphocytes, and T lymphocytes) were visualized on the transformed grid.

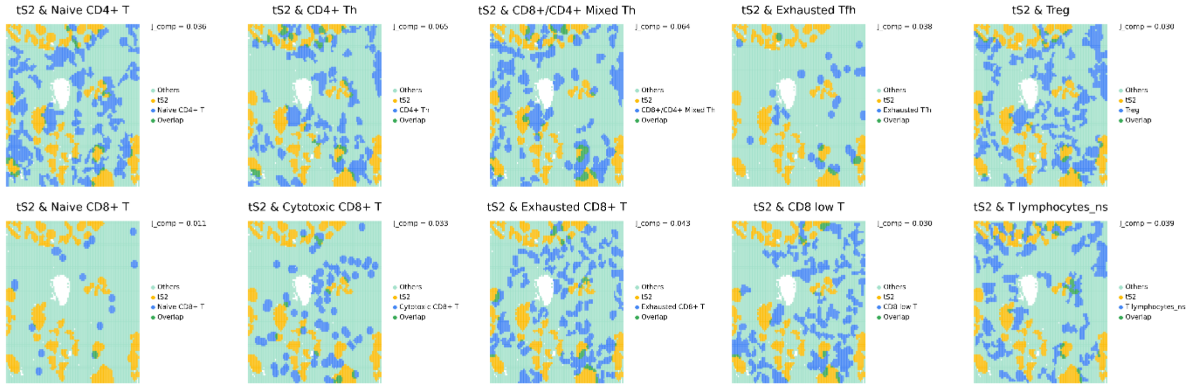


**Supplementary Fig. 10. The colocalization patterns between tS2 and multiple T cell subtypes in image-based SRT of lung cancer**

The key location of tS2 and multiple T cells subtypes were represented by CCs and the overlapping tissue domains were highlighted as the intersecting subregions between the two aggregated CCs.

**a**

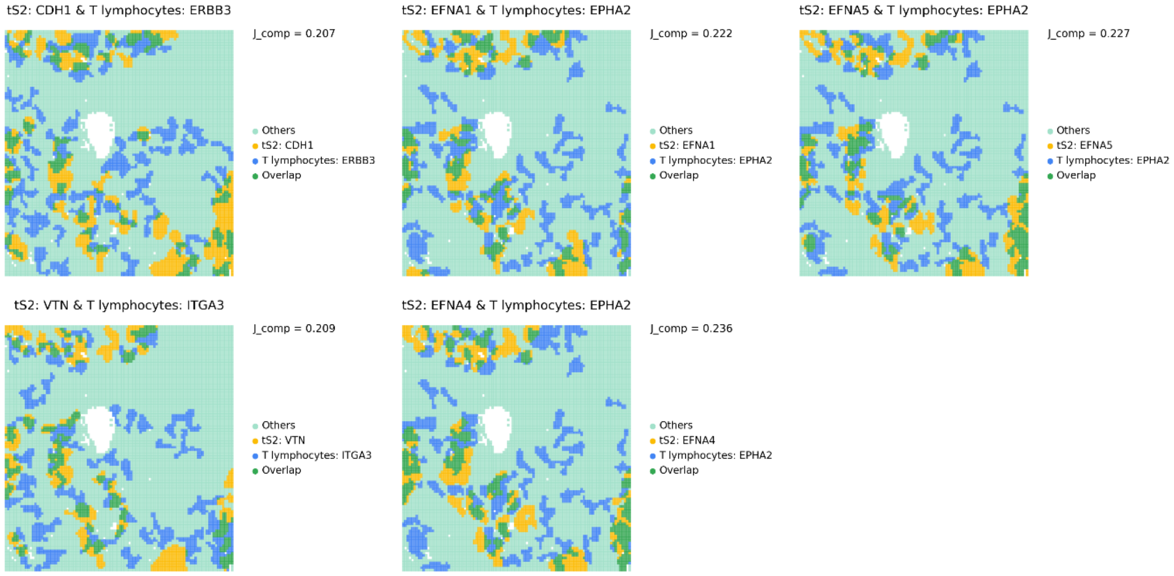
tS2: CDH1 & T lymphocytes: ERBB3
J_comp = 0.207
Others
tS2: CDH1
T lymphocytes: ERBB3
Overlap
tS2: EFNA1 & T lymphocytes: EPHA2
J_comp = 0.222
Others
tS2: EFNA1
T lymphocytes: EPHA2
Overlap
tS2: EFNA5 & T lymphocytes: EPHA2
J_comp = 0.227
Others
tS2: EFNA5
T lymphocytes: EPHA2
Overlap
tS2: VTN & T lymphocytes: ITGA3
J_comp = 0.209
Others
tS2: VTN
T lymphocytes: ITGA3
Overlap
tS2: EFNA4 & T lymphocytes: EPHA2
J_comp = 0.236
Others
tS2: EFNA4
T lymphocytes: EPHA2
Overlap

**b**

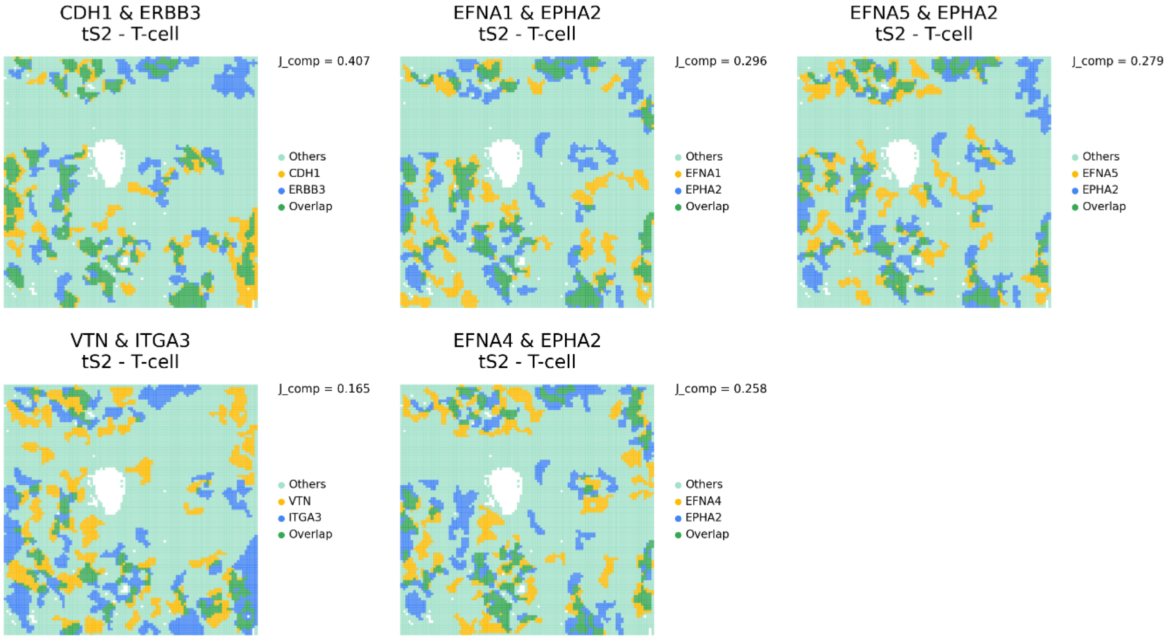
CDH1 & ERBB3
tS2 - T-cell
J_comp = 0.407
Others
CDH1
ERBB3
Overlap
EFNA1 & EPHA2
tS2 - T-cell
J_comp = 0.296
Others
EFNA1
EPHA2
Overlap
EFNA5 & EPHA2
tS2 - T-cell
J_comp = 0.279
Others
EFNA5
EPHA2
Overlap
VTN & ITGA3
tS2 - T-cell
J_comp = 0.165
Others
VTN
ITGA3
Overlap
EFNA4 & EPHA2
tS2 - T-cell
J_comp = 0.258
Others
EFNA4
EPHA2
Overlap

**Supplementary Fig. 11. Validation of a method to estimate cell type-specific LR interaction in barcode-based SRT.**

To validate an approach to estimate cell type-specific LR interaction in barcode-based SRT, an image-based SRT dataset was utilized since it can provide real cell type-specific expression profiles. (a) First, the ligand expression in tS2 and receptor expression in T cells were applied to find the key niche of LR interactions. The results were considered a reference standard. (b) Then, the alternative approach suggested for barcode-based SRT was applied and the tS2 and T cell-specific LR interaction was predicted. It was predicted by intersecting regions between the two aggregated CCs of the given LR pairs and colocalized domain of tS2 and T cells. The location of modified CCs was mapped to the tissue and the overlapping tissue domain was considered the key niche for tS2 and T cell-specific LR interactions.

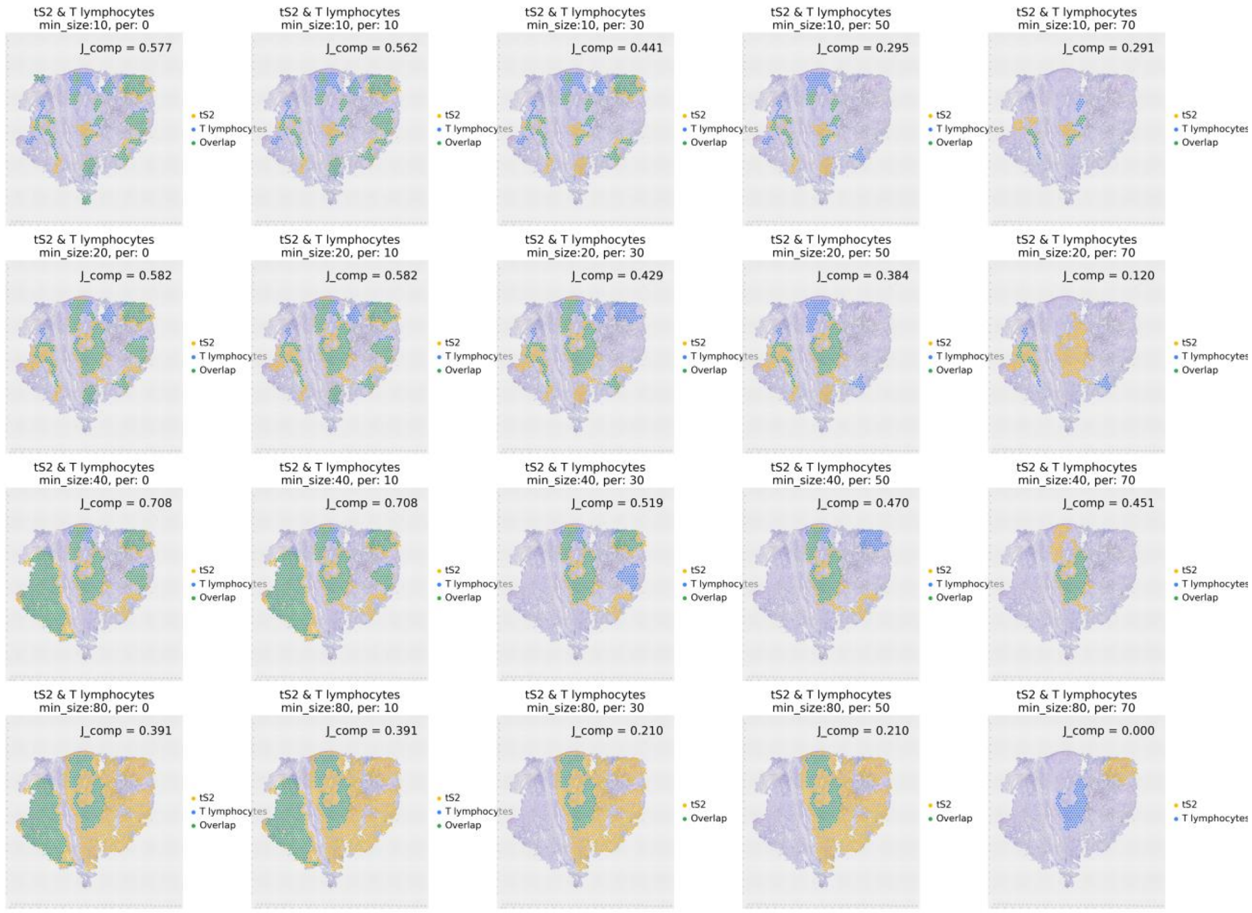


**Supplementary Fig. 12. The aggregated CCs obtained across various parameters in image-based SRT of lung cancer**

The aggregated CCs were calculated between tS2 and T cells by changing the two parameters: minimum size of CCs represented by the number of spots or grids and percentile threshold to remove the CCs with low average feature value. Minimum number of CCs were changed from 10 to 80 (10, 20, 40, 80) and percentile value from 0 to 70 (0, 10, 30, 50, 70).

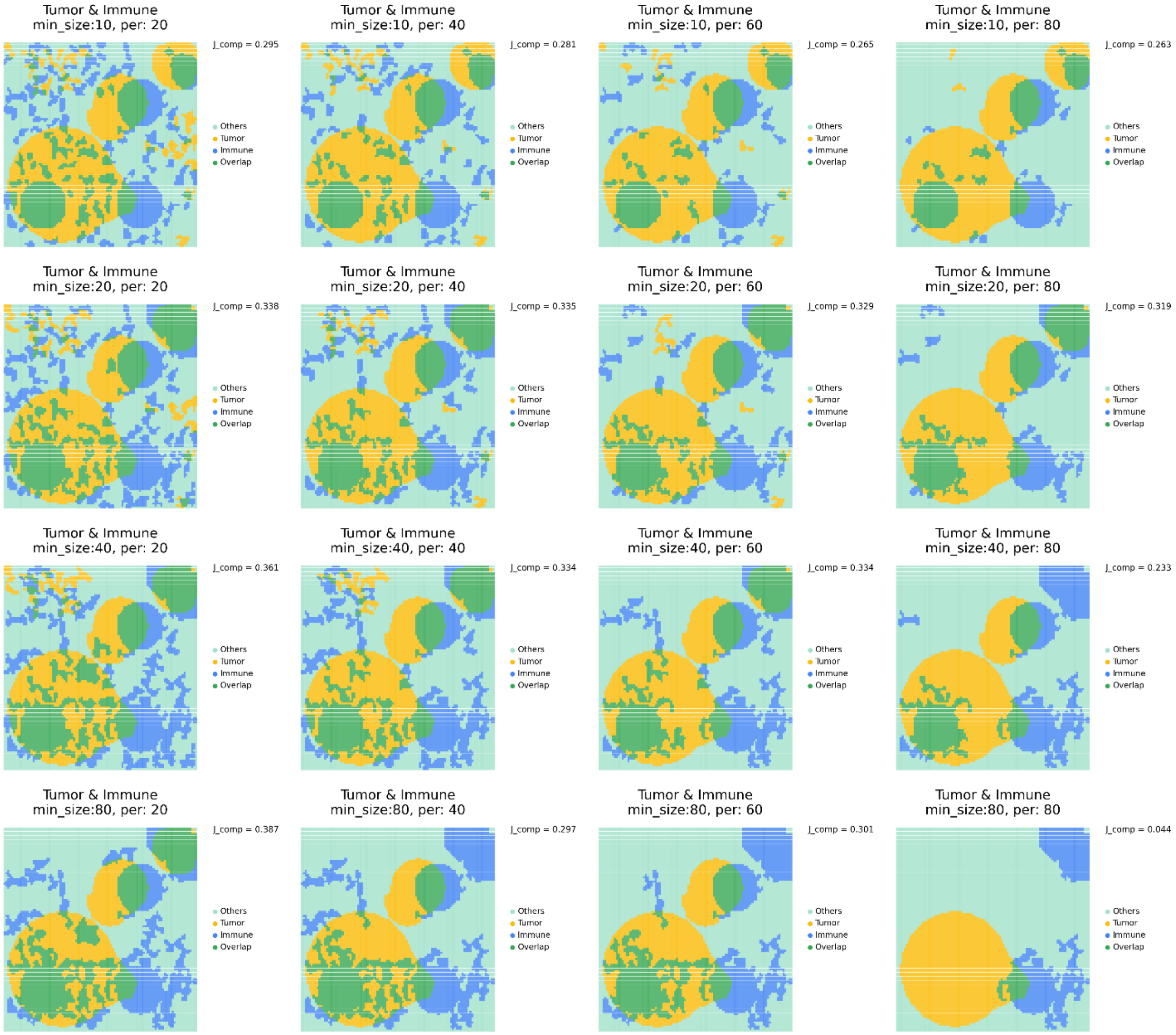


**Supplementary Fig. 13. The aggregated CCs obtained across various parameters in simulated dataset**

The aggregated CCs were calculated between tS2 and T cells by changing the two parameters: minimum size of CCs represented by the number of spots or grids and percentile threshold to remove the CCs with low average feature value. Minimum number of CCs were changed from 10 to 80 (10, 20, 40, 80) and percentile value from 20 to 80 (20, 40, 60, 80).
